# Unexpected mechanisms of sex-specific memory vulnerabilities to acute traumatic stress

**DOI:** 10.1101/2025.03.25.645300

**Authors:** Rachael E. Hokenson, Kiara L. Rodríguez-Acevedo, Yuncai Chen, Annabel K. Short, Sara A. Samrari, Brinda Devireddy, Brittany J. Jensen, Julia J. Winter, Christine M. Gall, Kiran K. Soma, Elizabeth A. Heller, Tallie Z. Baram

## Abstract

It is increasingly recognized that severe **acute** traumatic events (e.g., mass shooting, natural disasters) can provoke enduring memory disturbances, and these problems are more common in women. We probed the fundamental sex differences underlying memory vulnerability to acute traumatic stress (ATS), focusing on the role of the sex hormone, estrogen (17β-estradiol) and its receptor signaling in hippocampus. Surprisingly, **high** physiological hippocampal estrogen levels were required for ATS-induced episodic memory disruption and the concurrent sensitization and generalization of fear memories in both male and female mice. Pharmacological and transgenic approaches demonstrated signaling via estrogen receptor (ER)α in males and, in contrast, ERβ in females, as the mechanisms for these memory problems. Finally, identify distinct hippocampal chromatin states governed by sex and estrogen levels, which may confer an enduring vulnerability to post-traumatic memory disturbances in females.

## Main

With the increasing prevalence of mass shootings and climate-driven natural catastrophes, it is becoming evident that acute traumatic events (ATS) can provoke enduring disturbances of memory, and that these problems are more common in women. Persistent memory disturbances of several types occur in up to 30% of individuals exposed to earthquakes, the September 11 terrorist attacks or mass shootings^1–3^. These enduring consequences distinguish ATS from typical acute stresses, perhaps because of its severity, uncontrollability or complexity that includes concurrent physical, emotional and social elements^2,4–8^. Trauma-related memory disturbances commonly include deficits in episodic memory^2–4,9^ and a generalization of trauma-related memory cues^10^. When persistent and associated with intrusive trauma memories, these traumatic stress aftermaths are classified as post-traumatic stress disorder (PTSD)^11,12^. PTSD affects 6-8% of the population and exerts tremendous tolls on human potential and healthcare costs. PTSD is twice as common in women than men, even when controlling for trauma type^13–17^. The basis for these striking sex differences remains unknown and may relate to sex differences in brain organization, operations^18–26^ and vulnerability to stress^27–33^.

We previously discovered that ATS causes spatial memory disturbances in male mice and in female mice stressed during proestrus, when physiological estrogen levels are at the cyclical maximum. Surprisingly, memory in female mice stressed during estrus, the cyclical nadir of estrogen levels, was resilient to ATS^5–7^. Both male and female hippocampus synthesize estrogens and express estrogen receptors (ERs). Indeed, hippocampal levels of estrogens in males have been reported to approximate those in proestrous females^34^ in contrast to systemic levels that are always higher in females^34–36^. Hippocampal estrogen levels may govern the degree of activation of ERs, including ERα and ERβ that generally mediate the enduring effects of estrogens via direct or indirect interactions with the chromatin^24,37^.

The possibility that high physiological levels of estrogen may contribute to memory vulnerability to ATS, is unexpected, because estrogen generally promotes memory functions^38–44^ in both sexes^40,41^. However, the interaction of high levels of estrogen and acute traumatic stress is unclear^45–48^. Restoring estrogen to gonadectomized females protects memory from stress^43,49,50^. However, deleterious effects of high physiological levels of estrogen during stress have been reported^22,51,52^, suggesting a more complex relation of hormone levels, stress and memory.

Here we examined the memory disturbances following ATS to identify the basis of sex-specific vulnerabilities to this trauma and their underlying mechanisms. We identified an ATS-induced disruption of several types of memory in male and proestrous, but not estrous, female mice. In vulnerable mice, females were more sensitive to ATS than males, and their memory disturbances persisted. The use of both pharmacology and transgenic conditional hippocampal ER deletions demonstrated that ATS-provoked memory disturbances are mediated by ERα in males and ERβ in females. Finally, distinct sex- and estrogen-level dependent hippocampal chromatin states and expression of ER target genes associated with ATS outcomes, and may poise mice to vulnerability or resilience to ATS.

## Results

### Acute traumatic stress (ATS) disrupts hippocampus-dependent memory in a sex- and estrus cycle-dependent manner

Trauma-related memory disturbances commonly include deficits in episodic memory^2–4,9^. Here, we now demonstrate that ATS-induced spatial memory disturbances are persistent, and are associated with fear-memory generalization. Using a stress-free test for spatial memory, the object location memory (OLM) task in which mice preferentially explore the object moved to a novel location, we find that both male and proestrous female mice exposed to ATS and then trained in the task after a two-hour rest period did not preferentially explore the moved object^5,6^. In contrast, female mice in estrus, when hippocampal estrogen levels are lower, were resilient^7^. OLM performance remained poor in both vulnerable male and female mice also a week after ATS (Fig 1B).

**Fig. 1.**
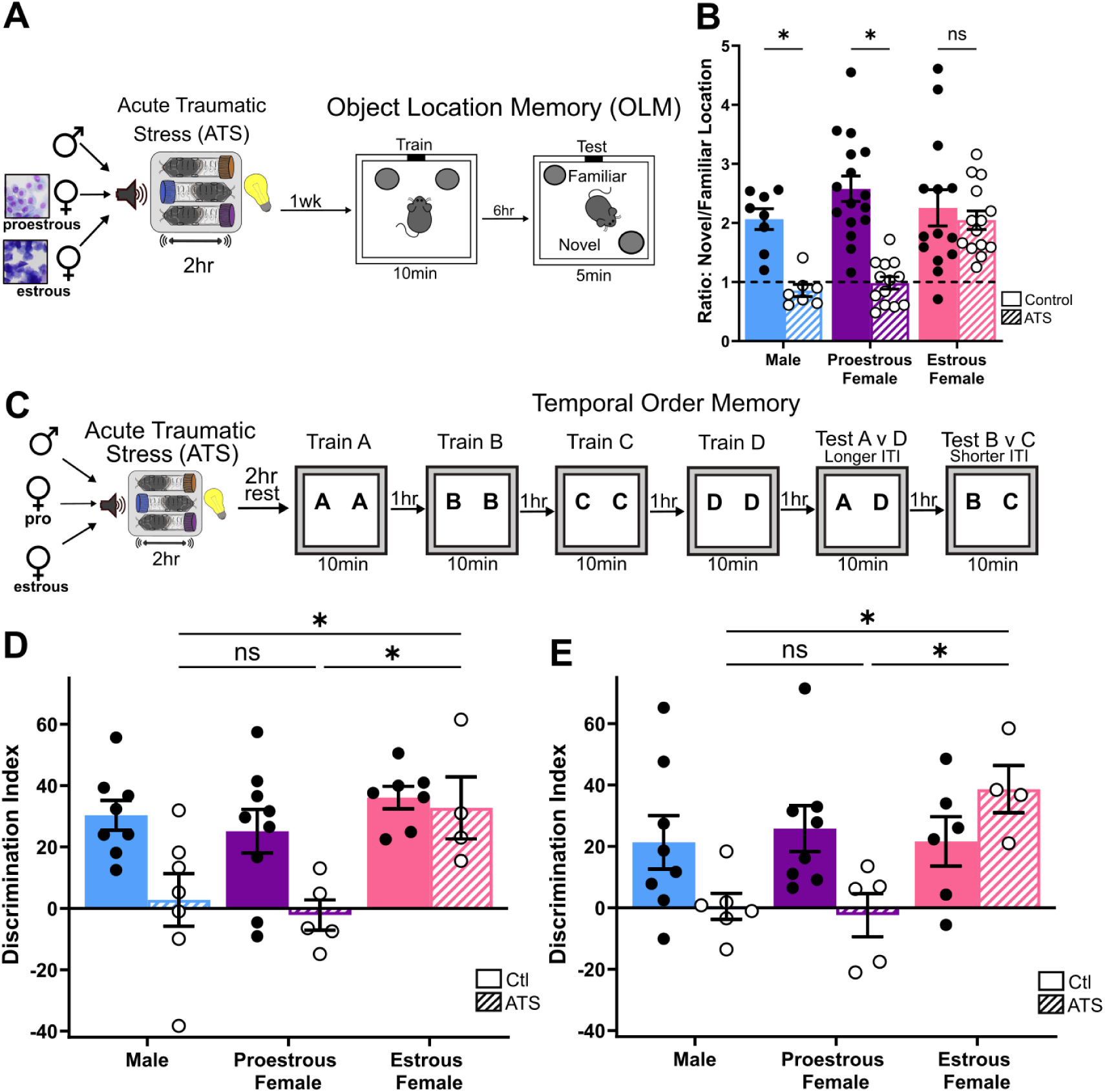
Acute traumatic stress disrupts hippocampus-dependent memory in male and proestrous female mice. (**A**) Male, proestrous, and estrous female mice were exposed to acute traumatic stress (ATS) and object location memory (OLM) was assessed one week later. Mice were trained with two identical objects for 10 minutes. One object was moved and mice were tested 6 hours later. (**B**) OLM was disrupted in male and proestrous female, but not estrous female mice (N = 7-16/group; 2-way ANOVA with Šidák’s multiple comparisons), (**C**) Male, proestrous, and estrous female mice were exposed to acute traumatic stress (ATS) and temporal order memory was assessed after a 2hr rest period. (**D**) Temporal order memory for objects A (remote) vs D (recent) was impaired by ATS in male and proestrous female mice. (N = 4-9/group; 2-way ANOVA with Šidák’s multiple comparisons). (**E**) Temporal order memory for objects B (remote) vs C (recent) was impaired by ATS in male and proestrous female mice. (N = 4-8/group; 2-way ANOVA with Šidák’s multiple comparisons). Each point represents an individual mouse. Bars display the mean ± SEM. Post-hoc comparisons are displayed above the graph. * = *P*<0.05, # = *P*<0.10.

Temporal order memory, a second type of episodic memory, was also impacted by ATS in a sex and hormone-level dependent manner. Control mice preferentially explored time-remote versus recent items with both long (3h, Fig 1D) and short (1h, Fig 1E) intervals between sessions. ATS disrupted temporal order memory in male and in proestrous female mice, whereas estrous females were again resilient (Fig 1D-E). These disruptions of object location- and temporal-memory could not be attributed to differences in exploration times during training or testing sessions (Fig S1).

OLM and temporal order are inherent, exploiting mice’s preference for novelty. We tested if <u>rewarded</u> spatial memory was also affected by ATS. In a contextual reward learning task (Fig S2A), control male and female mice learned the reward location of the apparatus and preferred the side paired with palatable food. ATS impaired contextual reward learning in male and proestrous females whereas estrous females remained resilient (Fig S2B). Poor contextual reward performance was not a result of reduced preference for the reward, as food consumption in ATS mice during conditioning was not reduced (Fig S2C).

Together, the above discoveries indicate that ATS provokes disturbances of several hippocampus-dependent memory types in male and proestrous female mice. In contrast, females stressed during estrus are resilient.

### ATS augments trauma-related memories in a sex- and estrus cycle-dependent manner

In humans, traumatic stress may lead to strong recurrent trauma memories and augment sensitivity to even partial cues of the stressful event (fear generalization,^11^). To test analogous sensitivity and generalization processes in mice, ATS was paired with an odor (almond) and an object (50ml tube): An almond-scented restraint tube was placed for an hour in home cages immediately following ATS. Control mice were exposed to the same cues in their home cage without ATS. We then tested mice for <u>avoidance</u> of concordant trauma cues (odor and object), partial cues (almond odor with a different object, or peppermint odor with the restraint tube) or neutral cues (neutral scent and object) (Fig 2A).

**Fig 2.**
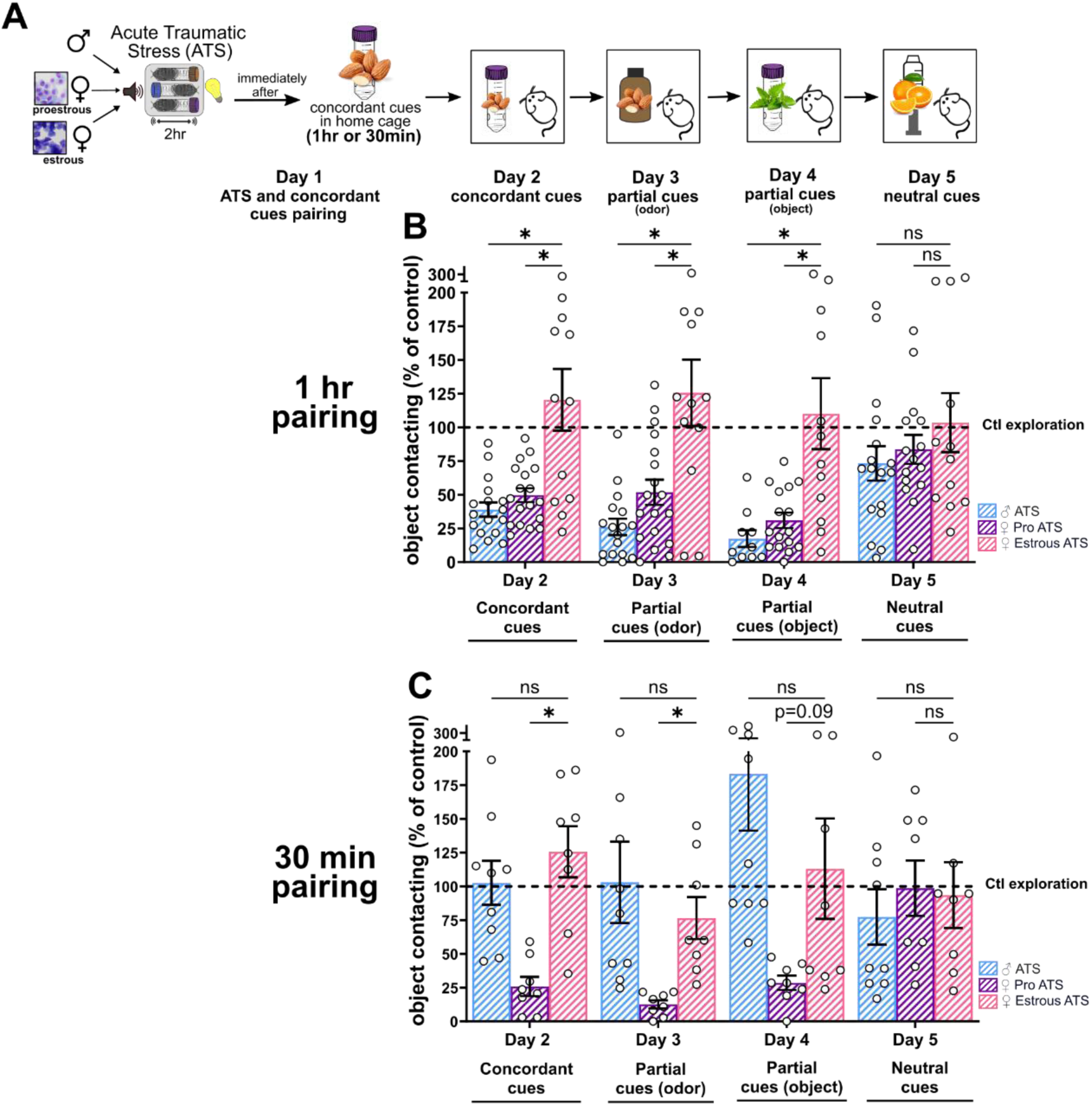
Acute traumatic stress provokes memory of and sensitivity to ATS-paired cues in a sex- and estrous cycle-dependent manner. (**A**) Male, proestrous, and estrous female mice were exposed to acute traumatic stress (ATS). Immediately after ATS, an almond scent (“ATS odor”) and a conical restraint tube (“ATS object”) were placed into the home cage for one hour or 30 minutes. Control mice were introduced to the cues in their home cage without ATS. Avoidance of concordant ATS cues, partial ATS cues, and neutral cues was assessed in the following weeks. (**B**) When pairing cues with ATS for one hour, male ATS and proestrous female ATS mice avoided concordant and partial ATS cues in the week following ATS compared to controls. Estrous female ATS mice did not avoid the ATS-related cues. All groups explored neutral cues equally (N = 10-18/group; Mixed-effects analysis with Dunnett’s multiple comparisons test). (**C**) In a separate cohort, ATS was paired with cues for 30 min. Proestrous female ATS mice avoided concordant and partial ATS cues compared to controls. Male and Estrous female ATS mice did not avoid the ATS-related cues. All groups explored neutral cues equally (N = 8-9/group; 2-way ANOVA with Dunnett’s multiple comparisons test). Each point represents an individual mouse. Bars display the mean ± SEM. Post-hoc comparisons are displayed above the graph. * = *P*<0.05, # = *P*<0.10

Male ATS mice and proestrous ATS females avoided the concordant trauma cues relative to controls (Day 2). However, they also avoided the partial cues, odor (Day 3) or object (Day 4), suggesting loss of specificity of the ATS memory (Fig 2B). In contrast, female mice stressed during estrus approached both concordant and partial ATS cues to the same degree as control females (Fig 2B, Table S1). All ATS mice explored neutral cues for the same duration as controls (Day 5), indicating that only cues associated with ATS were avoided.

The studies above identify a putative role for high physiological estrogen levels, but not sex, in the augmented avoidance of trauma-related cues. We reasoned that the apparent lack of sex-differences might be the result of an overwhelming exposure to the cue that masks important sex-dependent differences in sensitivity to ATS memories. To test this possibility, we modified the above experiment, shortening the duration of pairing of the cues (the almond-scented restraint tube) from 60 to 30 minutes. This approach revealed robust vulnerabilities of female mice: Even with the abbreviated pairing, proestrous ATS females continued to avoid both concordant and partial trauma cues (Days 2-4, Fig 2C, Table S1). In contrast, male ATS mice no longer avoided trauma-paired cues (Fig 2C). Exploration of neutral cues did not differ across groups (Day 5). These data indicate that proestrous female mice form an association of cues with ATS upon shorter pairing durations than males, i.e., they are more sensitive to ATS-related cue memories than males.

### Sex-dependent persistence of ATS-induced memory disturbances

In people, ATS may provoke enduring disturbances of memory that last for months and years, and these problems are more common in women^13^. Therefore, we probed both spatial and trauma-cue memories in mice during the weeks and months following ATS, to investigate their persistence and the potential underlying role of sex.

Object location memory (OLM) disturbances persisted at 1,4, and 8 weeks following ATS in vulnerable mice of both sexes (Fig 3A-C). However, robust sex differences in the sensitivity to enduring memory disturbances were uncovered by allowing mice to train longer (15 vs 10 minutes) at 4 or 8 weeks after ATS, together with shortening the interval to testing to 4 hours (Fig 3D). In this more lenient paradigm, spatial memory deficits persisted in proestrous female mice; whereas the more lenient task permitted ATS male mice to remember the location of a previously seen object (Fig 3E-F). These results indicate that ATS-provokes more intense memory disturbances in proestrous females than in males.

**Fig 3.**
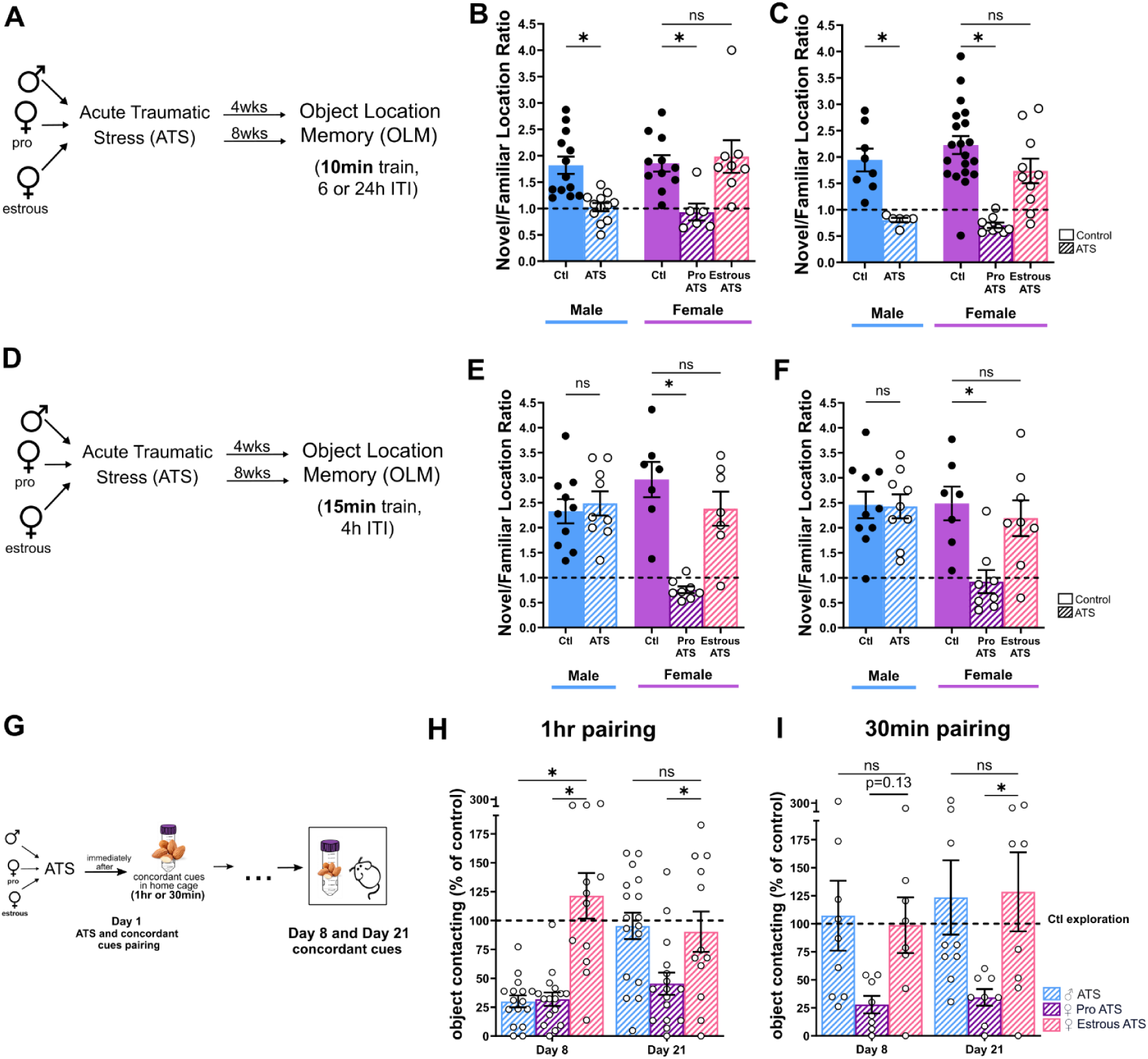
Acute traumatic stress provokes enduring memory disruptions in a sex-specific manner. (**A**) Male, proestrous, and estrous female mice were exposed to acute traumatic stress (ATS) and object location memory (OLM) was assessed 4 weeks or 8 weeks later. Mice were trained with two identical objects for 10 minutes. One object was moved and mice were tested 24 hours (for 4 week cohort) or 6 hours later (for 8 week cohort) later. (**B**) OLM was impaired in male and proestrous female, but not estrous female mice, four weeks following ATS (N = 6-13/group; Kruskal-Wallis with Dunn’s multiple comparisons), and (**C**) and eight weeks following ATS (N = 6-20/group; Kruskal-Wallis with Dunn’s multiple comparisons). (**D**) In a different cohort of mice, male, proestrous, and estrous female mice were exposed to acute traumatic stress (ATS) and object location memory (OLM) was assessed 4 or 8 weeks later. Mice were trained for 15 minutes with two identical objects. One object was moved and mice were tested 4 hours later. OLM was impaired by ATS in proestrous female but not male or estrous female at (**E**) 4 weeks and (**F**) 8 weeks after ATS (N = 7-10/group, Ordinary one-way ANOVA with Šidák’s multiple comparisons). (**G**) Male, proestrous, and estrous female mice were exposed to acute traumatic stress (ATS). Immediately after ATS, an almond scent and a conical restraint tube were paired to ATS by introducing these cues for one hour or 30 minutes in the home cage. Control mice were introduced to the cues in their home cage without ATS. (**H**) With 1 hour pairing, male ATS and proestrous female ATS mice avoided concordant cues on day 8 while on day 21 only proestrous female ATS mice avoided concordant trauma cues (N = 10-16/group; Mixed-effects analysis with Dunnett’s multiple comparisons test). (**I**) With 30 minutes of pairing, proestrous female ATS mice avoided concordant cues on day 8 and day 21 while male ATS and estrous ATS mice did not avoid trauma cues (N = 8-9/group; 2-way ANOVA with Dunnett’s multiple comparisons test). Each point represents an individual mouse. Bars display the mean ± SEM. Post-hoc comparisons are displayed above the graph. * = *P*<0.05, # = *P*<0.10

Augmented vulnerability of females to the enduring impact of ATS involved trauma-cue memories as well (Fig 3G). With a one hour cue pairing, both male and proestrous female mice tested a week after ATS (day 8), avoided concordant cues. In contrast, 3 weeks after ATS (day 21), while proestrous females continued to avoid concordant cues, ATS males explored the cues at control levels (Fig 3H). In addition, when cue pairing was reduced to 30 min, male ATS mice no longer avoided concordant trauma cues at either 1 or 3weeks later, whereas ATS proestrous females remained sensitive (Fig 3I).

Together, the findings above indicate that female sex confers a heightened sensitivity to ATS, with augmented severity and persistence ATS-induced memory disturbances.

### Hippocampal estrogen synthesis and signaling via estrogen receptors is required for ATS-induced spatial memory disturbances

The studies above, demonstrating vulnerability to ATS-induced memory problems in females *only* during estrous cycle phases characterized by high hippocampal estrogen levels^53^, and in males, in which hippocampal estrogen levels are reportedly high^34^, raised the possibility that high physiological levels of the key estrogen, 17β-estradiol, may be required for ATS-induced memory deficits. To test this idea, we first measured hippocampal estrogen levels using an ultrasensitive and specific liquid chromatography tandem mass spectrometry assay (Fig. 4A ^54,55^). Male levels of hippocampal 17β-estradiol (33.43 ± 5.50 pg/g) were comparable to those of proestrous females (41.88 ± 6.61 pg/g), and higher than those of estrous female mice (19.89 ± 2.19 pg/g).

**Fig 4.**
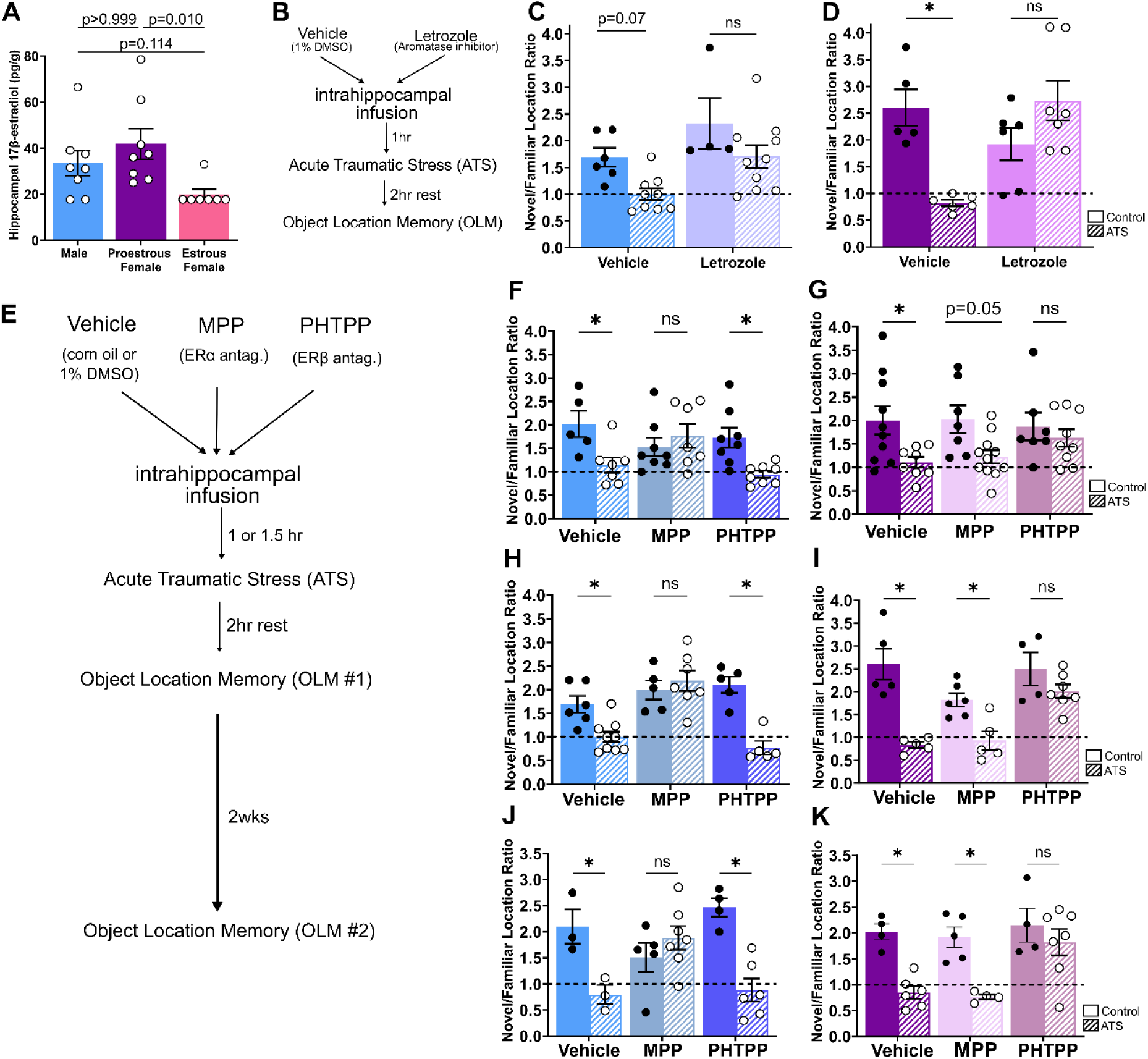
Spatial memory was protected from acute traumatic stress by pharmacologically blocking estradiol production. These effects are mediated by estrogen receptor alpha in males and estrogen receptor beta in females. (**A**) Hippocampal estrogen (17β-estradiol) levels are higher in mice vulnerable to ATS (male and proestrous female) compared to mice resilient to ATS (estrous female). (N = 7-8/group, one-way ANOVA). Most estrous female values are below the lower limit of quantification (25 pg/g). (**B**) Male and female mice were implanted with indwelling, bilateral cannula aimed at the dorsal hippocampus. One hour before ATS, vehicle (1% DMSO in saline) or an aromatase inhibitor (6 or 60ng letrozole) were infused. 2 hours after ATS, mice were tested on the object location memory task. (**C**) Object location memory was protected from ATS in male mice treated with letrozole (N = 4-10/group). (**D**) Likewise, object location memory was protected from ATS in proestrous female mice treated with letrozole (N = 5-7/group). (**D**) Male and proestrous female mice were given subcutaneous vehicle (corn oil), an estrogen receptor alpha antagonist (MPP, 0.5mg/kg), or an estrogen receptor beta antagonist (PHTPP, 0.5mg/kg) 1.5hr prior to acute traumatic stress (ATS). Object location memory (OLM) was assessed after a 2hr rest. Alternatively, male and proestrous female mice with indwelling bilateral cannulae aimed at dorsal hippocampus were given infusions of vehicle (1% DMSO in saline), an estrogen receptor alpha antagonist (MPP, 0.05pmol/hemisphere), or an estrogen receptor beta antagonist (PHTPP, 0.1pmol/hemisphere) 1hr prior to acute traumatic stress (ATS). Object location memory (OLM) was assessed after a 2hr rest. (**F**) OLM was disrupted by ATS in vehicle and PHTPP treated male mice. However, ATS did not disrupt OLM in MPP treated male mice (N = 5-8/group) (**G**) OLM was disrupted by ATS in vehicle and MPP treated proestrous female mice. However, ATS did not disrupt OLM in PHTPP treated proestrous female mice (N = 7-11/group). (**H**) In males, blocking ERα with MPP (0.05pmol/hemisphere) infused in hippocampus, protected memory from ATS (N = 5-9/group).(**I**) In proestrous females, blocking hippocampal ERβ with PHTPP protected memory from ATS (N = 4-7/group). Two weeks later, mice who received intrahippocampal infusions of vehicle or ER blockers completed a second object location task. (**J**) In males, mice infused with vehicle or PHTPP had lasting ATS-induced memory disruptions, while mice with ERα blocked during ATS (MPP infusion) remained protected (N = 3-7/group). (**K**) In females, mice infused with vehicle or MPP and stressed during proestrus had lasting memory deficits, while mice with ERβ blocked during ATS (PHTPP infusion) remained protected (N = 4-7/group). Each point represents an individual mouse. Bars display the mean ± SEM. All behavior data are analyzed with 2-way ANOVA with Šidák’s multiple comparisons. Post-hoc comparisons are displayed above the graph. * = *P*<0.05.

Hippocampal estrogens are derived from local synthesis as well as systemic sources^35,56,57^. We tested if acute local inhibition of hippocampal estrogen synthesis protects mice from ATS-induced memory problems. We infused the aromatase inhibitor letrozole into the dorsal hippocampus. Similar to chronic systemic aromatase inhibition^58^, acute intrahippocampal infusion of an aromatase inhibitor protected male and proestrous female mice from ATS-induced memory disturbances (Fig. 4B-D), identifying a key role of hippocampal estrogen synthesis in mediating memory disruptions by stress.

How might high physiological levels of estrogens contribute to ATS-induced memory disturbances? Estrogen acts via several types of receptors and signaling cascades^43,59–63^, and the enduring actions of the hormone are thought to involve the direct or indirect interaction of the canonical ERs, ERα and ERβ, with the chromatin. To identify which ERs mediate ATS-induced spatial memory disturbances in male and female mice, we systemically blocked ERα and ERβ using a selective ERα antagonist (MPP, 0.5mg/kg), a selective ERβ antagonist (PHTPP, 0.5mg/kg), or vehicle (corn oil, sc) 90 minutes before ATS. In males, blocking ERα, but not ERβ, protected OLM in ATS mice with little influence on controls. In females, instead, only blocking ERβ rescued OLM from ATS-induced impairments (Fig 4E-G). We pinpointed the actions of the ERs to the hippocampus by administering MPP (0.05pmol/hemisphere), PHTPP (0.1pmol/hemisphere), or vehicle (1% DMSO in saline) directly to dorsal hippocampus 60 min prior to ATS. In accord with the systemic interventions, blocking hippocampal ERα selectively protected spatial memory in ATS males, and blocking ERβ protected vulnerable females (Fig 4E,H-I). These protections were enduring: while OLM remained impaired in vehicle-treated males or proestrous females tested two weeks after ATS, it remained intact in males with ERα blocked and in proestrous females with ERβ blocked during ATS (Fig. 4J-K). The aromatase inhibitor and ER blockers had minimal effect in controls, and their protection of ATS mice was not a result of altered object investigation (Fig S4). Thus, a transient, sex-specific block of hippocampal ERs during ATS suffices to provide lasting protection of memory.

### Focal hippocampal deletion of specific estrogen receptors in adult male and female mice protects from ATS-provoked memory disturbances in a sex-dependent manner

The pharmacological approaches described above supported a sex-specific role of distinct ERs in ATS-induced memory disturbances. To definitively test these unexpected results, we generated mice with a conditional deletion of hippocampal ERs, i.e., mice lacking ER alpha (ERαKO), ER beta (ERβKO), or both (ERαβKO) primarily in hippocampal principal cells (Fig 5A; Fig.S5A). We used a CamKinaseIIa dependent-cre-expressing mouse line, in which cre expression was largely confined to the hippocampal formation (Fig. S5A), leading to pronounced reduction of ER expression in hippocampus, but not, for example, in salient hypothalamic regions (Fig. 5B,C). We then employed a within-subject design to determine the role of each ER in ATS-induced memory disturbances First, male and female ER intact, ERαKO, ERβKO, or ERαβKO mice performed the OLM task. Then, a month later, the same mice were exposed to ATS (female mice in proestrus) and then tested again for OLM (Fig 5D). Prior to ATS, all the transgenic mice performed the task similarly to wild-type mice^5,7^. ATS disrupted OLM in ER intact male mice, and deletion of hippocampal ERα (ERαKO) or both receptors (ERαβKO) but not of ERβ (ERβKO) protected memory from ATS (Fig 5E). In contrast, in females, ATS disrupted OLM in ER intact proestrous mice, and deletion of hippocampal ERα did not protect hippocampus-dependent memory, whereas ERβKO and ERαβKO female mice performed the OLM task equally well before and after ATS (Fig 5F). As a control, memory in ER intact (Cre^-^) male and proestrous female mice was impaired by ATS regardless of the floxed ER genotype (Fig S5C-D). In the transgenic mice, as in the wildtype mice, the effects of ATS on memory and the protection by sex-specific deletion of ERs endured for at least 3 weeks after ATS (Fig S5B,E-H). Again, we excluded the possibility that memory disturbances or protection from these disturbances in ERKO mice resulted from differential object investigation during training or testing sessions (Fig S6).

**Fig 5.**
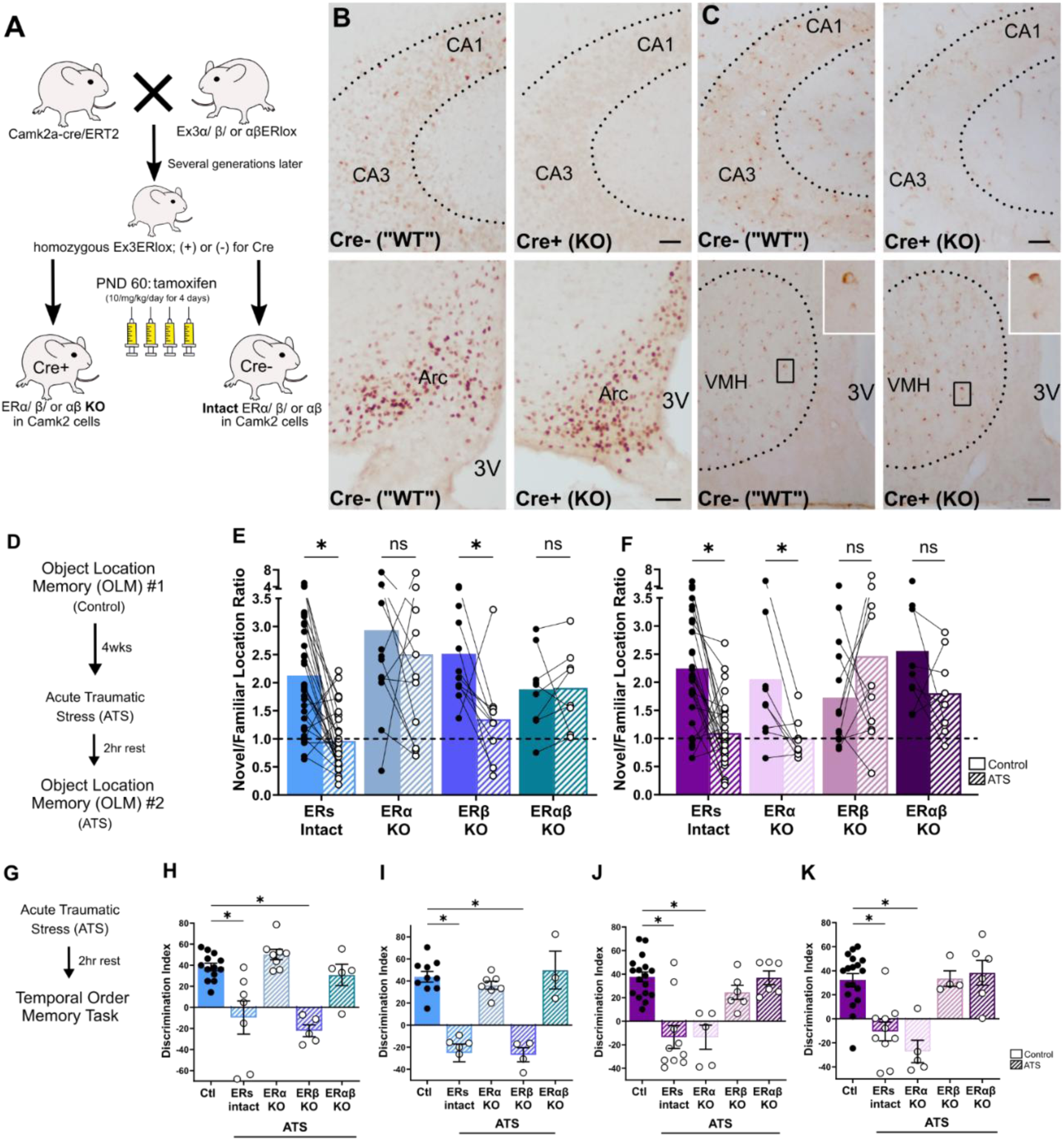
Focal hippocampal deletion of specific estrogen receptors in adult male and female mice protects from ATS-provoked spatial and temporal memory disturbances in a sex-dependent manner. (**A**) Hippocampal estrogen receptor knock-out (ERKO) mice were generated by crossing CamKII-Cre-ERT2 with mice with floxed estrogen receptor alpha, beta, or both alpha and beta. Resulting offspring were Cre^+^ (KO) or Cre^-^ (ERs intact) and homozygous for the floxed ER of choice. Mice were genotyped on postnatal day 14 and treated with tamoxifen (10mg/kg/day for 4 days) starting at PND 60. All behavior tasks were performed one month after cessation of tamoxifen treatment. (**B**) ERα levels in Cre^+^ floxed ERα mice (ERαKO) (top right) were reduced in hippocampus compared to Cre^-^ floxed ERα mice (ER intact) (top left). In contrast, ERα staining did not differ between Cre^+^ or Cre^-^ floxed ERα mice in the arcuate nucleus (Arc) (bottom). Scale bar = 50 µm. (**C**) Likewise, ERβ levels in ERβKO mice (top right) were reduced in hippocampus compared to ER intact mice (top left). In contrast, ERβ staining did not differ between ER intact and ERβKO mice in the ventromedial nucleus of the hypothalamus (VMH) (bottom). Scale bar = 50 µm. 3V = 3^rd^ ventricle. (**D**) Male and female mice of each ERKO genotype were tested on the object location memory task (control). One month later, mice were exposed to ATS then OLM was tested again after a 2 hour rest. (**E**) OLM was disrupted by ATS in ER intact and ERβKO male mice. However, ATS did not disrupt OLM in ERαKO and ERαβKO male mice (N = 8-39/group, Mixed-effects analysis with Šidák’s multiple comparisons). (**F**) OLM was disrupted by ATS in ER intact and ERαKO proestrous female mice. However, ATS did not disrupt OLM in ERβKO and ERαβKO proestrous female mice (N = 9-32/group, Mixed-effects analysis with Šidák’s multiple comparisons). (**G**) Male and proestrous female ERKO mice were exposed to acute traumatic stress (ATS) and temporal order memory was assessed after a 2hr rest period. In male mice, temporal order memory was disrupted in ER intact and ERβKO mice, but ERαKO and ERαβKO had intact memory in (**H**) Test A versus D (N = 3-8/group, ordinary one-way ANOVA, Šidák’s multiple comparisons) as well as (**I**) Test B versus C (N = 3-6/group, ordinary one-way ANOVA, Šidák’s multiple comparisons). In proestrous female mice, temporal order memory was disrupted in ER intact and ERαKO mice, but ERβKO and ERαβKO had intact memory in (**J**) Test A versus D (N = 4-11/group, ordinary one-way ANOVA, Šidák’s multiple comparisons) as well as (**K**) Test B versus C (N = 3-12/group, ordinary one-way ANOVA, Šidák’s multiple comparisons). Each point represents an individual mouse with connected points representing the same mouse at two time points. Bars display the mean ± SEM. Post-hoc comparisons are displayed above the graph. * = *P*<0.05.

The sex-specific roles of ERs in ATS-induced memory disturbances extended to temporal order memory (Fig 5G). Specifically, compared to controls, ATS disrupted temporal order memory for objects A vs D (Fig 5H) and objects B vs C (Fig 5I) in ER intact as well as ERβKO male mice, but not ERαKO and ERαβKO males (Fig 5H-I). In proestrous females, ATS disrupted temporal order memory for objects A vs D (Fig 5J) and objects B vs C (Fig 5K) in ER intact as well as ERαKO mice, but ERβKO and ERαβKO female mice were protected (Fig 5J-K). Again, there were no discernible differences by group in object investigation during training or testing sessions (Fig S7).

Together, the use of transgenic mice buttresses the pharmacological data and establishes sex-specific roles for hippocampal ERs in ATS-induced disturbances of spatial and temporal order memory, both components of episodic memory. Specifically, they demonstrate that ERα in males and ERβ in proestrous females mediate the impact of ATS on episodic memory.

### Focal deletion of sex-specific estrogen receptors prevents ATS-induced over-sensitivity and generalization of trauma cues

Having established the sex-specific role of ERα and ERβ in ATS-induced disturbances of spatial and temporal memory, we proceeded to investigate the role of the receptors in the augmented sensitivity to ATS-related cues using the selective deletion of ERs. We exposed vulnerable male and proestrous female mice with intact ERs (Cre negative) and those lacking ERα, ERβ or both receptors to ATS paired with a “trauma-odor” and “trauma object”, as described for the wild-type mice. Avoidance of concordant trauma cues as well as partial and neutral cues was then tested (Fig 6A). Male ATS mice with intact ERs avoided concordant and partial trauma cues (Days 2-4, Day 8, Table S2) relative to controls, in accord with wild-type mice (Fig 2B). Male mice lacking hippocampal ERα or both receptors, but not ERβKO, explored the trauma cues for the same duration as controls (Fig 6B). In proestrous females, hippocampal deletion of ERβ or both receptors, but not ERα alone, led female ATS mice to explore trauma cues for the same amount of time as controls (Fig 6C). In all genotypes and sexes, exploration of neutral cues did not differ across groups (Day 5).

**Fig 6.**
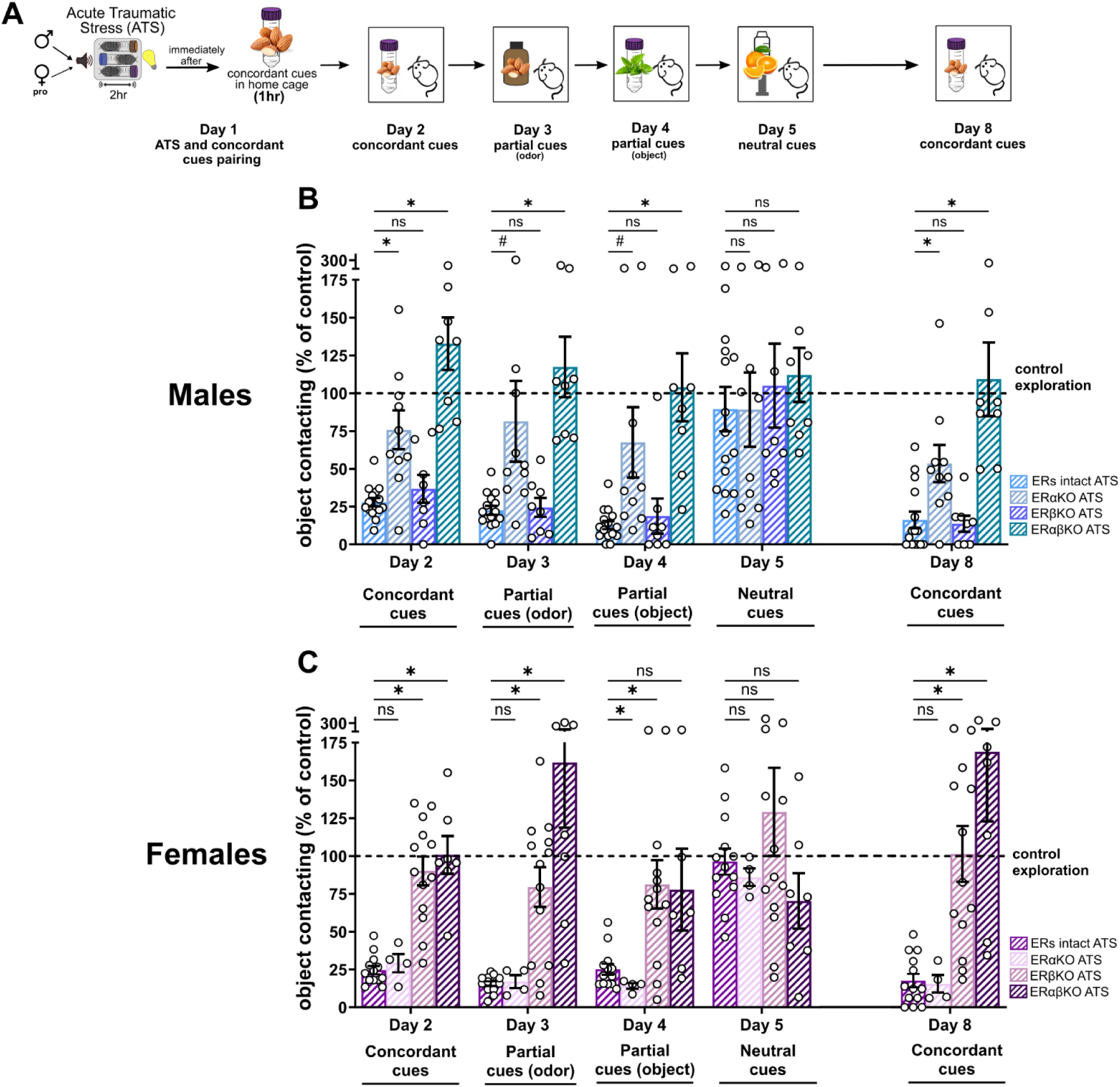
Focal deletion of hippocampal estrogen receptors prevented ATS-induced memory and generalization of trauma cues. (**A**) Male and proestrous female mice of each ERKO genotype were exposed to acute traumatic stress (ATS). Immediately after ATS, an almond scent and conical restraint tube cue were placed in the home cage for one hour. Avoidance of concordant trauma cues, partial trauma cues, and neutral cues was assessed in the following week. (**B**) ER intact and ERβKO Male ATS mice avoided concordant and partial trauma cues while ERαKO and ERαβKO male ATS mice approached the cues similar to control mice (N = 8-15/group, Mixed-effects analysis with Dunnett’s multiple comparisons test). (**C**) ER intact and ERαKO proestrous female ATS mice avoided concordant and partial trauma cues while ERβKO and ERαβKO proestrous female ATS mice approached the cues similar to control mice (N = 4-13/group, Mixed-effects analysis with Dunnett’s multiple comparisons test). Each point represents an individual mouse. Bars display the mean ± SEM. Post-hoc comparisons are displayed above the graph. * = *P*<0.05, # = *P*<0.10

Together, these data indicate that (a) ERs mediate ATS-provoked augmented sensitivity and ‘generalization’ of trauma cues in a sex-dependent manner and (b) the same ERs mediate ATS effects on the different types of memory disturbances in a given sex.

### ATS-induced memory disruption may derive from distinct hippocampal chromatin states at ER-associated TF motifs, governed by sex and estrogen levels

What might be the mechanisms for involvement of ERα in males and ERβ in females in mediating ATS-induced memory disruptions during high hippocampal estrogen states? A selective recruitment of each receptor type, driven by permissive or repressed chromatin states and by hormone levels, provides a plausible mechanism. Mouse hippocampus chromatin structure differs as a function of sex and estrous cycle phase^37^ such that ERα chromatin binding sites are differentially enriched in open chromatin in proestrous vs. (di)estrous females^21^. ERs regulate gene expression, in part, by recruiting histone modifying enzymes and transcription factors that promote or repress gene expression^64,65^. Such alterations of gene expression are intrinsic to memory processes and their disruption in disease^66–70^. Therefore, we reasoned that a basis of the sex- and hormone-level specific ATS vulnerability might derive from the effects of sex and hormone levels on chromatin states.

To test this possibility, we performed genome-wide chromatin profiling of active (enriched in H3K4me3), repressed (H3K27me3), and bivalent (harboring both H3K4me3 and H3K27me3) gene promoters in naïve proestrous and estrous females and in male mouse hippocampi (Fig. 7A). The results were striking: ATS resilient estrous females exhibited very few (44) unique permissive gene promoters, and, instead, 2307 unique repressive or bivalent gene promoters compared to ATS susceptible male and proestrous female groups (1415 permissive, 51 repressive or bivalent; Fig. 7B). In addition, a large number of permissive promoters was shared between proestrous females and males but not estrous females (Fig. 7C). In contrast, estrous females exhibited a distinct chromatin landscape, with a greater number of unique repressed (N = 265; Fig. 7D) and bivalent (N = 2,042; Fig. 7E) gene promoters.

**Fig 7.**
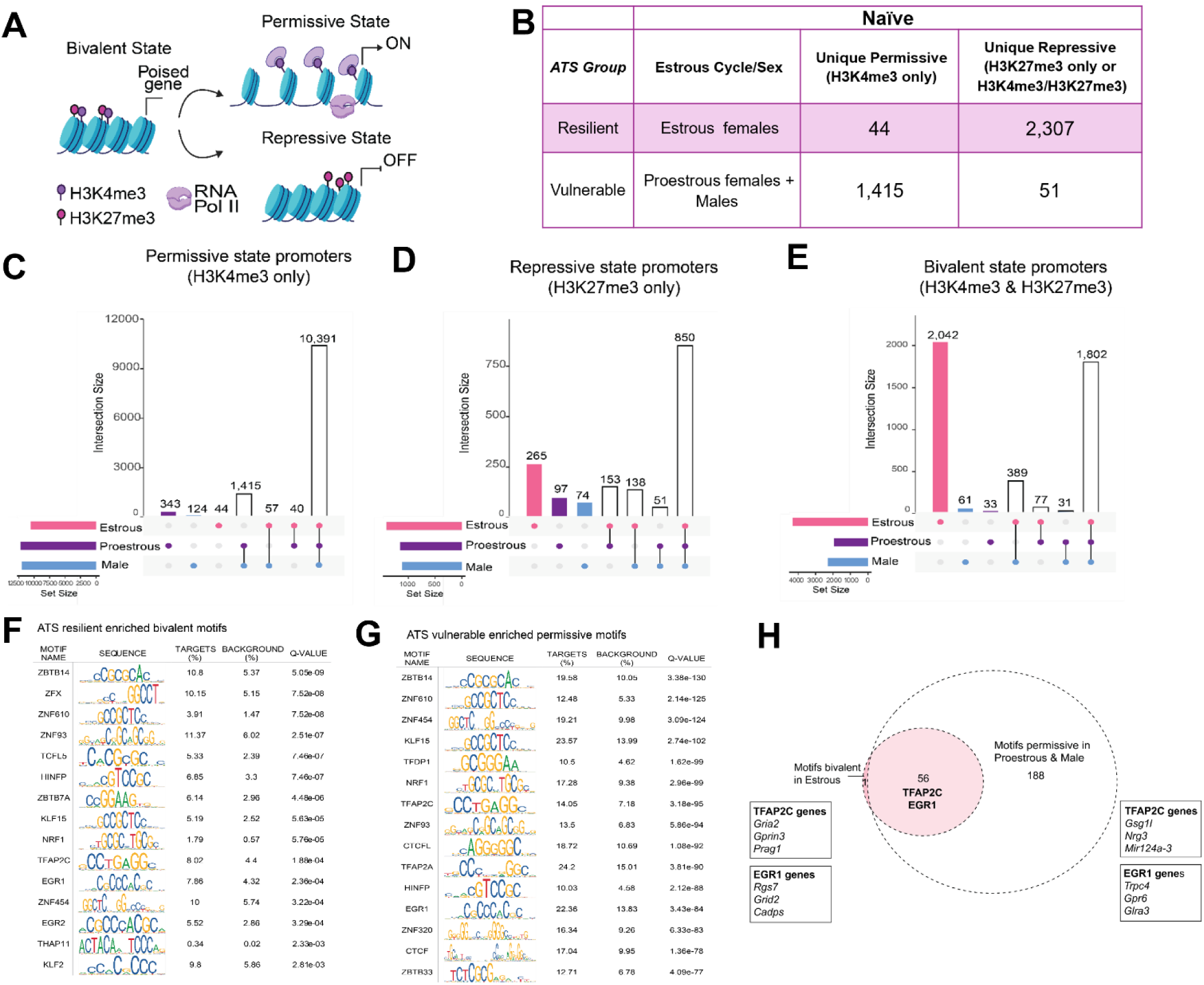
Sex and estrous cycle phase impact hippocampal chromatin state. (**A**) Bivalent genes possess both H3K4me3 and H3K27me3 histone posttranslational modifications. These genes are thus poised for transcriptional activation or repression. Upon an appropriate stimulus, loss of H3K27me3 promotes a permissive state (H3K4me3 only), while loss of H3K4me3 reinforces a repressive state (H3K27me3 only). Chromatin state was analyzed in naïve (stress-free) male, proestrous female, and estrous female mouse hippocampus. (**B**) Summary counts of unique gene promoter regions (+/-100bp from TSS) enriched in permissive or repressive chromatin in naïve ATS-vulnerable or resilient mice. ATS-resilient females show higher bivalent and repressive states while males and proestrous females show higher permissive states. (**C**) Permissive state promoters (H3K4me3 only) were more numerous in male and proestrous females than estrous females. (**D**) In parallel, more repressive state promoters (H3K27me3 only) were identified in estrous females than males and proestrous females. (**E**) Promoter bivalency (both K3K4me3 and K3K27me3) was more frequent in estrous females. (**F**) Top 15 enriched motifs including ER-associated Egr1 and TFAP2 family of transcription factors were bivalent in estrous females and (**G**) permissive in proestrous females and males. (**H**) Summary motif figure showing that transcription factor motifs that are enriched in bivalent chromatin unique to estrous females are instead permissive in proestrous females and males (Motif-containing genes differ by groups).

The predominance of bivalent chromatin states in estrous females prompted us to hypothesize that specific estrogen-related transcription factors (TFs) within this unique gene set may be differentially activated in ATS resistant versus resilient mice. To test this possibility, we performed Simple Enrichment Analysis (SEA) to identify TF motifs within genes that were bivalent uniquely in estrous females. We identified a strong enrichment of TFAP2 motifs, including TFAP2C / Estrogen Receptor Factor 1 (ERF1), a target of both ERα and ERβ^71^ (Fig. 7F,H). We also found enrichment of Egr1 motifs in estrous-female-specific bivalent chromatin, consistent with ATAC-seq data showing Egr1 motifs are more accessible in proestrous compared to estrous female hippocampus^37^. These findings suggest that whereas ER-interacting TFs are enriched in active / permissive promoters in ATS-vulnerable mice, they are within bivalent genes in ATS resilient females, and thus may not support ATS-induced activation of ER target genes (Fig. 7G,H).

To determine the potential consequences of the differences in chromatin states on ER-dependent gene expression in estrous females and the two vulnerable groups, we performed gene expression profiling using RNA-sequencing of the same samples profiled by chromatin state^73^. Bivalent genes (marked by both H3K4me3 and H3K27me3) were minimally expressed, as expected^72^ whereas permissive (H3K4me3-marked) chromatin correlated with high transcript expression, and repressive H3K27me3 correlated with lower expression (Fig. 8A,S8). Indeed, global gene expression profiles were sex- and estrogen-level specific by principal component analysis: PC1 and PC2 explained 67% of the variance (Fig.8B), and the ATS-vulnerable groups clustered together. The number of differentially expressed genes (DEGs) was greatest between estrous females and either proestrous females (N = 243) or males (N = 139), and was lowest between the two ATS-vulnerable groups (N = 27, Fig 8C). These transcriptional data, in accord with the chromatin states, suggest that ATS vulnerable mice share a similar transcriptional profile that differs from ATS resilient females. Gene ontology (GO) analyses identified synaptic signaling and organization leading the functions of DEGs between estrous and proestrous females (Fig. 8D), while metabolic processes were enriched in the estrous versus male comparison (Fig. 8E). No significant GO terms distinguished DEGs between ATS-vulnerable proestrous female and males.

**Fig. 8.**
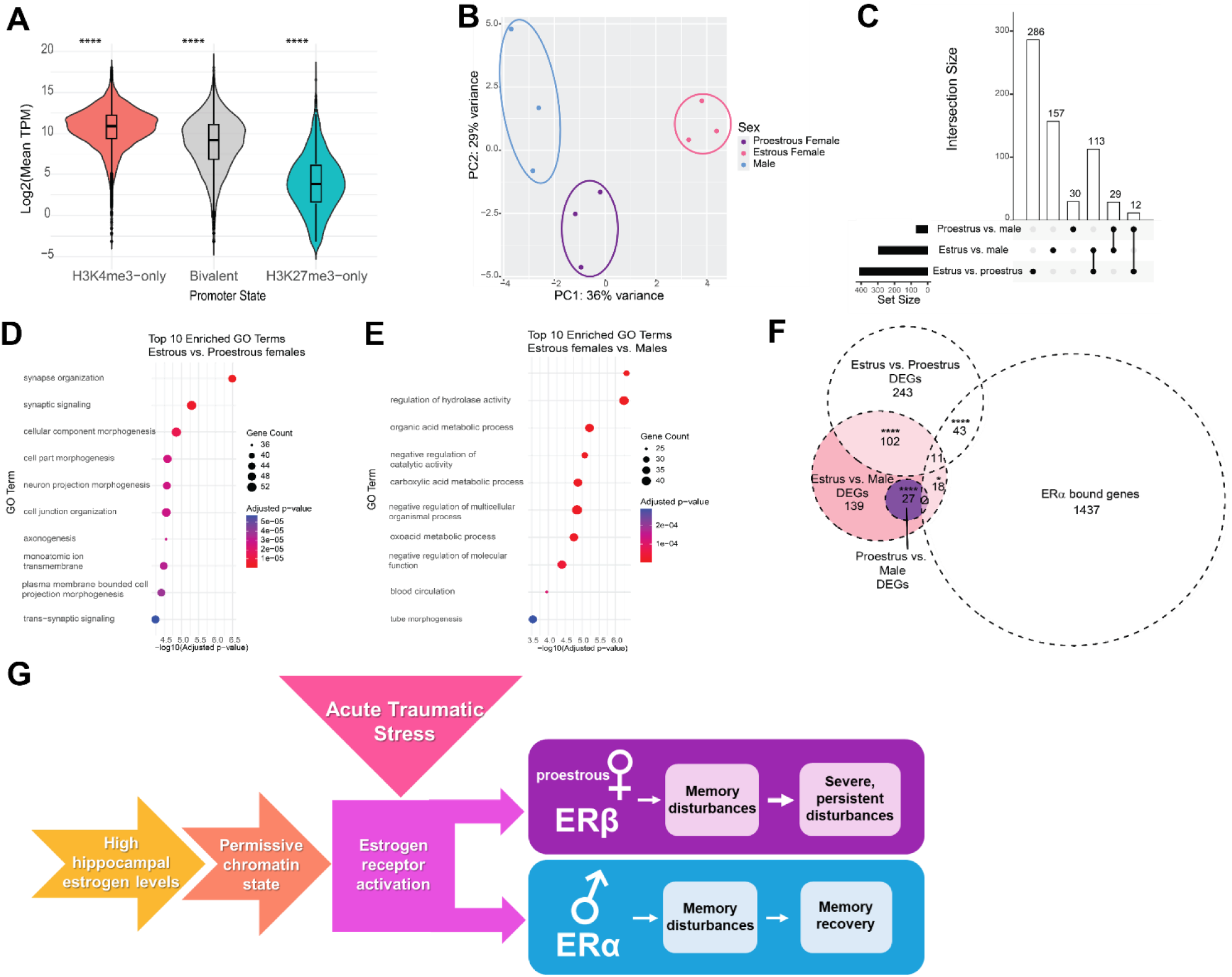
Differential gene expression by sex and estrous cycle phase in mouse hippocampus. (**A**) Gene expression levels across different promoter chromatin states. Promoters (±100 bp from TSS) were categorized as H3K4me3-only, H3K27me3-only, or bivalent (H3K4me3 + H3K27me3) and intersected with RNA-seq expression data. Violin plots show that genes marked by H3K4me3-only (N = 11,057) have significantly higher TPM compared to bivalent (N = 3,999) and H3K27me3-only (N = 1,328) promoters (Wilcoxon test, adjusted p < 0.05 for all comparisons). (**B**) Principal component plot for RNA-seq data of male, proestrous, or estrous female mouse hippocampus. The three biological replicates per sex and estrous cycle phase (N = 3/group) show consistent clustering. Males and proestrous females cluster closer together on PC1. (**C**) Upset plot portraying unique and overlapping differential expressed genes in all comparisons. (**D**) Gene ontology enrichment analysis shows top 10 enriched GO terms, with some relating to synaptic signaling pathways in the estrous vs. proestrous female comparison (**E**) whereas top 10 enriched GO terms for the estrous female vs. male comparison are metabolic-related terms. (**F**) Overlap analysis of DEGs in male, proestrous female, and estrous female hippocampus and ERα targets (targets identified using dataset described in ^24^ shows significant overlap by Fisher’s exact test of the estrous vs. proestrous and the estrous vs. male DEGs. (**G**) A cartoon depicting the proposed mechanisms by which ATS provokes memory disturbances, governed by sex and hippocampal estrogen levels, and mediated by sex-specific estrogen receptors (ERs). Proestrous female mice were more sensitive to ATS-induced memory deficits and these deficits were more persistent than those in males.

Finally, we aligned the DEGs with established targets of ERα^24^, to probe for potential effects of sex and estrogen levels on these transcriptional profiles. We identified 43 putative ERα targets differentially expressed between estrous and proestrous females, consistent with differential regulation driven by the hormone. In contrast, only 2 DEGs between proestrous females and males were targets of ERα (Fig 8F).

Taken together, our discoveries support a model (Fig. 8G) in which hippocampal estrogen levels contribute to chromatin states and gene expression programs in naïve males and females that predispose groups with high-physiological estrogen to vulnerability to ATS. Sex may govern the activation of distinct gene expression programs and the longevity of the ATS-induced memory disruption via differential involvement of the two major ERs.

## Discussion

Understanding the basis of sex-specific vulnerabilities to acute traumatic events (ATS), a growing global risk-factor for memory disturbances including PTSD, is crucial for potential preventative and mitigation strategies. Here we probed these fundamental sex differences, focusing on hippocampal estrogen levels and receptor signaling. We established that high physiological hippocampal estrogen levels are permissive to ATS-induced disruptions of several memory types. We then demonstrated, using both genetic and pharmacological approaches, estrogen receptor (ER)α in males and, in contrast, ERβ in females as the mediators of these memory problems. We further identified sex-and hormone-level dependent hippocampal chromatin states that enable receptor-mediated gene expression and consequent signaling cascades, leading to vulnerability or resilience to ATS.

In accord with the enduring nature of trauma-induced memory disturbances in people, we established the persistence of ATS-induced memory disturbances in mice, which lasted over two months. In addition, ATS-experiencing mice exhibited measures of generalization of trauma-related cues that resemble fear-generalization, a core symptom of PTSD in the human. As with the spatial memory problems, the trauma-cue generalization took place in males and proestrous females, whereas, in contrast, females in estrus were resilient. Indeed, the levels of hippocampal estrogen *during the ATS* determined vulnerability while estrus-cycle dependent estrogen level variation at the time of testing was inconsequential. Thus, the current studies recapitulate crucial elements of post-traumatic memory disruption, providing a robust framework for deciphering the underlying mechanisms.

Here, we measured hippocampal estradiol levels by LC-MS/MS and demonstrated that high physiological levels of hippocampal estrogen are required for vulnerability to ATS as both lower levels during estrus and the inhibition of hippocampal estrogen synthesis protected males and females from ATS-induced memory disturbances. This was unexpected because estrogen generally enhances memory^39–43^. and when estrogen is reduced surgically or pharmacologically, its replenishment protects from stress^43,49,50^. These strong data contrast with other work showing that high physiological estrogen levels during stress may exacerbate its impacts^22,51,52^. A way to reconcile these ostensibly inconsistent results is to conceptualize a non-linear relation of hippocampal estrogen levels, stress and memory. For example, estrogen may modulate memory in an “inverted U” manner, as shown for the stress hormone corticosterone^74–77^.

In addition to the role of high physiological hippocampal estrogen levels, sex governs the vulnerability to ATS-induced memory disturbances: Varying the severity of the ATS experience (duration of the trauma cue pairing) or the conditions for assessing memory disturbances unmasked augmented vulnerabilities in females compared with males with similar hippocampal estrogen levels, suggesting additional sources of the sex-specific female vulnerability to ATS. Whereas major sex-differences exist in the operation of memory systems, ranging from synaptic to circuit levels^26,41^, an additional mechanism stems from the role of ERβ rather than ERα signaling in female ATS-induced memory deficits. Target genes of the two receptors differ, so that ERβ-specific signaling cascades may lead to augmented sensitivity to ATS. A role of ERβ in the effect of stress in females is supported by several studies. For examples, ERβ in central amygdala regulates the consequences of repeated social stress in female rats^78^, and a sex-specific ERβ usage in females has been identified in heroin-conditioned extinction memory^79^.

The basis of sex- and hormone-level specific signaling cascades that enable ATS-induced memory deficits has not been previously elucidated. Here we examined hippocampal chromatin states and gene expression in male and proestrous and estrous females. Hippocampal gene promoters unique to ATS-resilient estrous females were largely in a repressed or bivalent chromatin state while those unique to ATS-vulnerable proestrous female and male mice were largely permissive. Notably, these differences included ER target genes: A preponderance of unique ER target genes in naïve ATS-vulnerable groups were poised for expression upon a suitable stimulus, including ATS (Fig.7G,H), potentially executing ATS-induced memory disturbances. In contrast, the same genes were repressed in ATS-resilient estrous females. One example is estrogen-sensitive TFAP2C transcription factor^71^ :TFAP motifs were abundant in bivalent genes in estrous females and, in contrast, in permissive genes in proestrous females and males. Thus, this estrogen-sensitive TF may differentially regulate expression of numerous genes in these groups.

Our work identifies robust, sex-dependent roles of ERα and ERβ in ATS-induced memory disturbances, but does not address the potential role of the G-protein coupled estrogen receptor (GPER1). GPER1, expressed in hippocampus^43,80^, mediates several estrogen effects on cognitive functions, yet its contribution to ATS-induced memory problems remains unknown. In addition, estrogen and sex may influence levels or activity of corticotropin releasing hormone^81,82^ or glucocorticoid receptors^83^, both of which play roles in stress-related memory problems^84,85^.

In conclusion, we establish important sex differences in memory vulnerabilities to acute traumatic stress, a growing global problem. We demonstrate an unexpected permissive role of high physiological hippocampal estrogen levels to ATS-induced memory disturbances in both males and females, and identify sex-and hormone dependent chromatin states as a potential contributor. Combining pharmacological and transgenetic approaches, we identify the sex-specific role of ERα and ERβ in ATS-induced disturbances of memory, suggesting the sex-dependent sensitivity and longevity in proestrous females relate to their use of ERβ, the brain-targets of which are as yet unknown. Because of the evident vulnerability of women to cognitive and emotional consequences of stress^86–89^ uncovering the mechanisms involved will advance our ability to address these important issues in women’s health^90^.

## Supplemental Figures

**Fig. S1.**
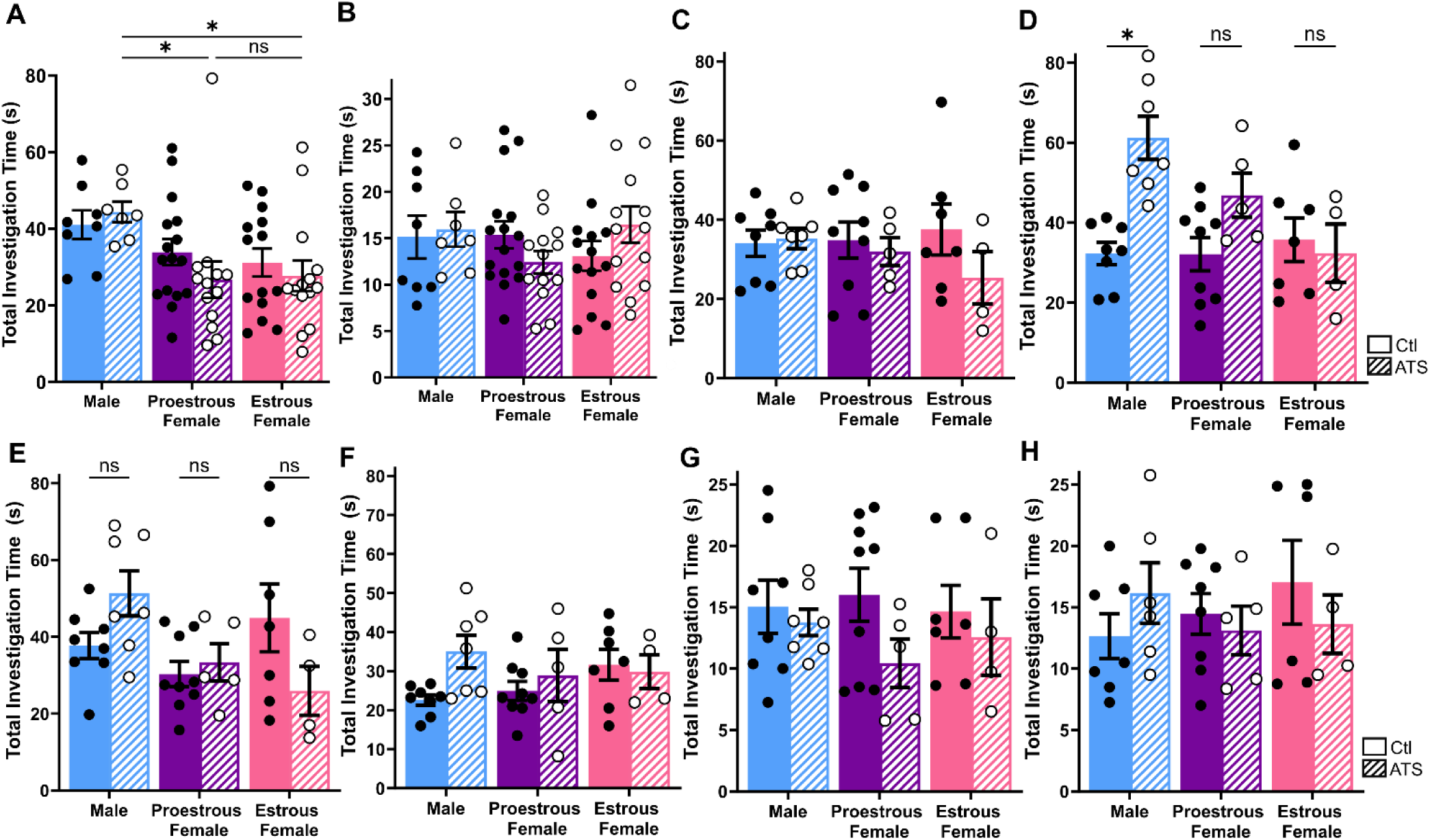
Disrupted object location memory or temporal order memory by acute traumatic stress were not a result of differences in investigation times during training or testing. In the object location memory task conducted 1 week after ATS, total object investigation time (**A**) during OLM training differed by sex/cycle, but not by ATS, (**B**) and there were no group differences in investigation during testing (N = 8-16/group, 2-way ANOVA). In the temporal order memory task, there were no group differences in exploration in (**C**) Object A training. (**D**) During object B training, ATS did impact investigation time but, rather than decreasing exploration, ATS drove an increase of exploration in males. (**E**) For total object investigation time during object C training, there was an interaction of sex/cycle and ATS on investigation time but post-tests were not significant. There were no group differences in total object exploration during (**F**) object D training, (**G**) object A vs D testing, or (**H**) object A vs D testing (N = 4-9/group, 2-way ANOVA). Each point represents an individual mouse. Bars display the mean ± SEM. Post-hoc comparisons are displayed above the graph. * = *P*<0.05, # = *P*<0.10

**Fig. S2.**
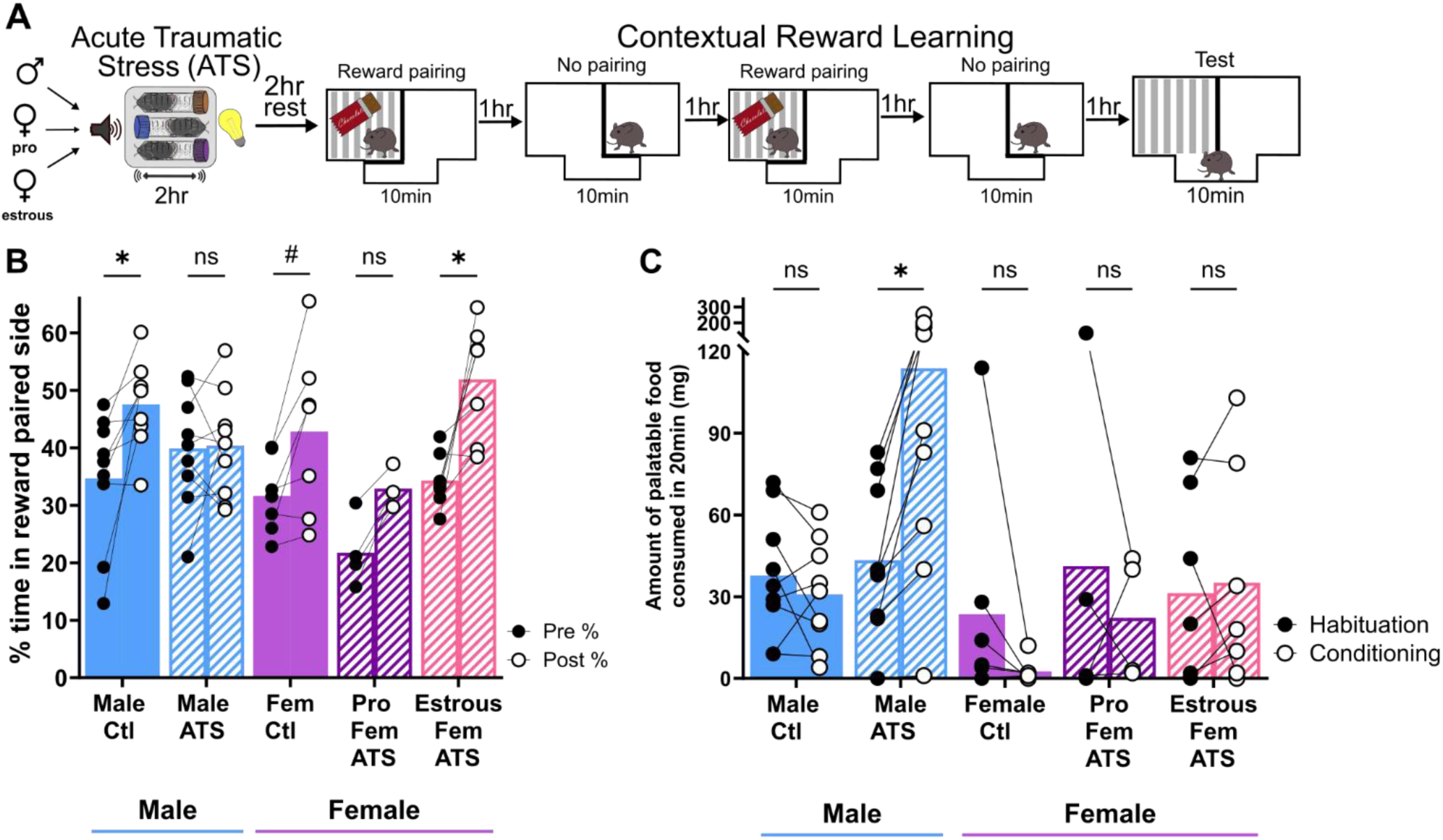
Contextual reward learning is disrupted by ATS in a sex and estrous cycle dependent manner. (**A**) Mice were habituated to an apparatus with two distinct chambers and acclimated to palatable food (cocoa pebbles) prior to acute traumatic stress (ATS). Male, proestrous, and estrous female mice were exposed to ATS. One side of the apparatus was paired with cocoa pebbles while the other side was not paired with any stimulus. Following conditioning sessions, mice were returned to the open chamber to assess a change of side preference. (**B**). Percent of time spent in the reward paired side was increased from habituation to the test phase in control male, control female, and female estrous ATS mice. Male ATS and proestrous female ATS did not increase time spent in the reward paired side after conditioning (N = 4-9/group, 2-way repeated measures ANOVA with Dunnet’s multiple comparisons) (**C**) ATS-induced disruption of reward learning was likely not due to reduced interest in consuming as reward, as amount of palatable food consumed during the conditioning sessions was not reduced compared to habituation. Indeed, ATS drove male mice to consume more palatable food (N = 4-9/group, 2-way repeated measures ANOVA with Dunnet’s multiple comparisons). Each point represents an individual mouse. Bars display the mean. Post-hoc comparisons are displayed above the graph. * = *P*<0.05, # = *P*<0.10

**Fig. S3.**
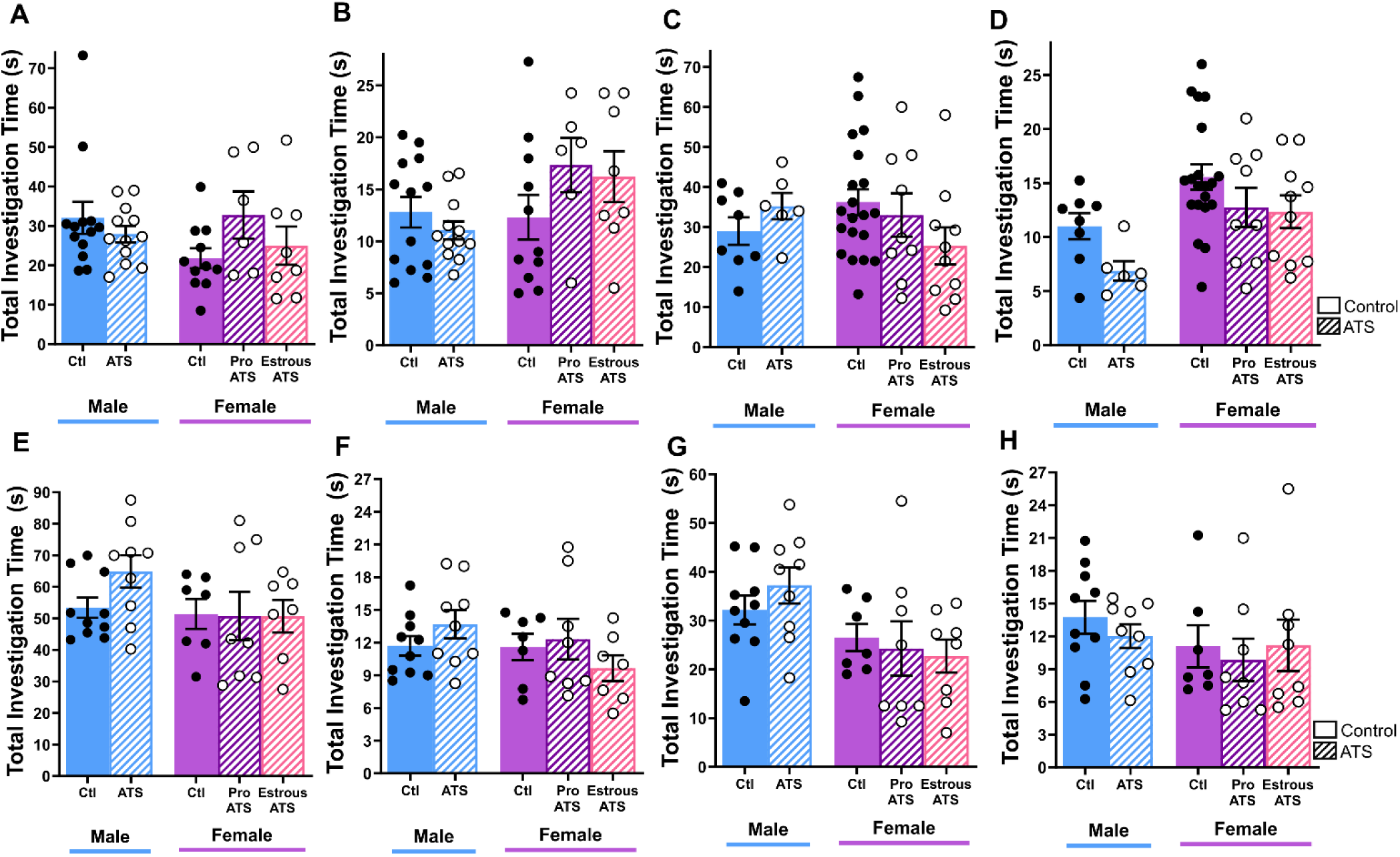
Enduring, sex-specific disruptions of object location memory by acute traumatic stress were not a result of a difference in investigation times during training or testing. Male, proestrous, and estrous female mice were exposed to ATS then tested on an object location memory test 4 or 8 weeks later. Mice were trained for 10 minutes then tested 6 or 24 hours later. 4 weeks after ATS, total object investigation did not differ between groups during object location (**A**) training or (**B**) testing (N = 6-13/group, Kruskal-Wallis). 8 weeks after ATS, total object investigation time did not differ between groups during (**C**) training but (**D**) there was a difference in investigation time during testing, though this was not driven by ATS (N = 6-20/group, Kruskal-Wallis). In separate cohorts, object location memory training was conducted for 15 minutes, and testing occurred four hours later. 4 weeks after ATS, there were no group differences in object exploration during (**E**) training or (**F**) testing (N = 7-10/group, ordinary one-way ANOVA). 8 weeks after ATS there were no group differences in object exploration during (**G**) training or (**H**) testing (N = 7-10/group, ordinary one-way ANOVA). Each point represents an individual mouse. Bars display the mean ± SEM. Post-hoc comparisons are displayed above the graph. * = *P*<0.05, # = *P*<0.10

**Fig. S4.**
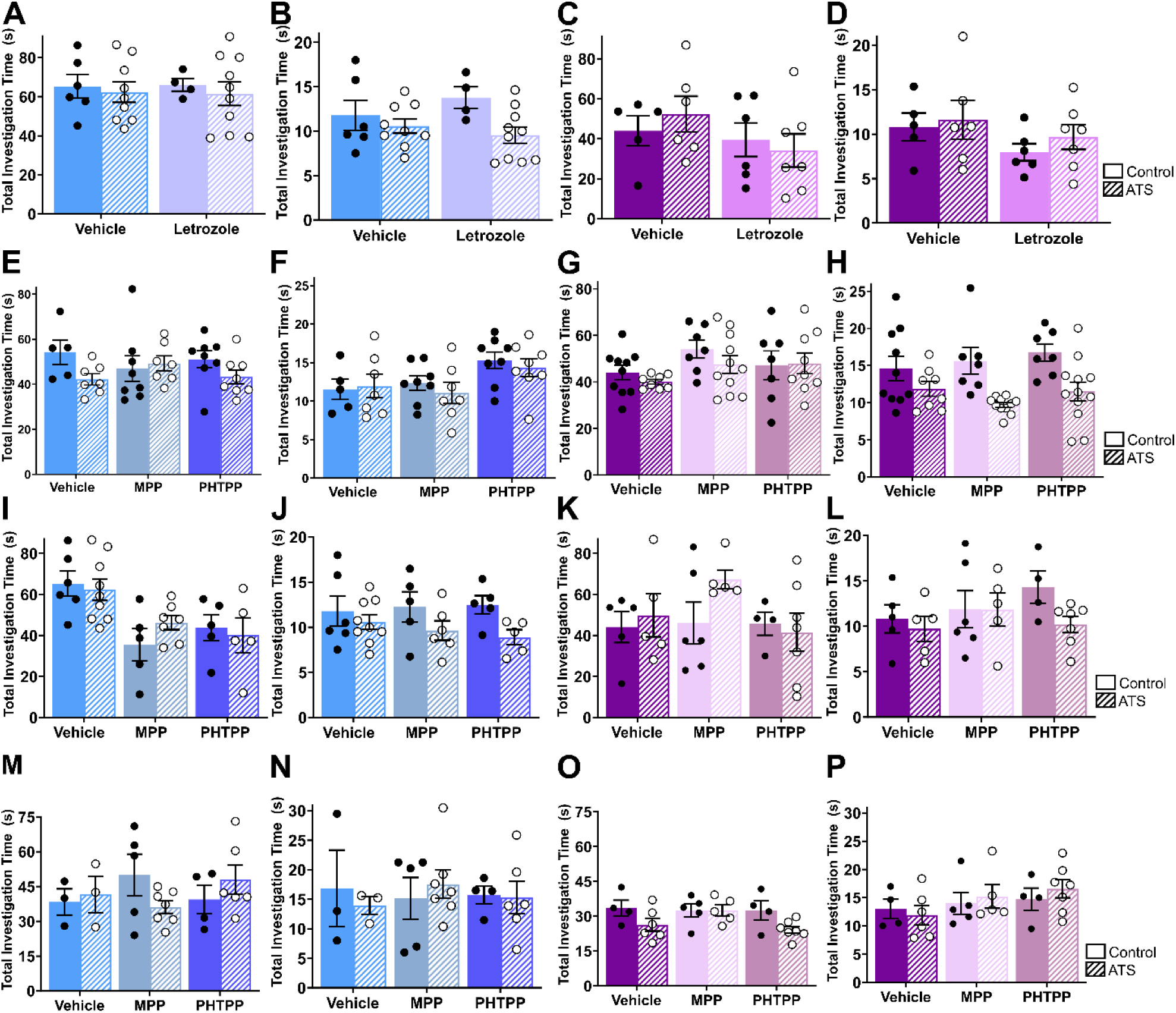
Spatial memory disruptions and protection with aromatase inhibition or estrogen receptor antagonism were not a result of a difference in investigation times during training or testing. In mice infused with the aromatase inhibitor letrozole, there were no differences in object investigation in the OLM task during (**A**) training in males (**B**) testing in males (N = 4-10/group, 2-way ANOVA), (**C**) training in females, (**D**) or testing in females (N = 5-7/group, 2-way ANOVA). When treated subcutaneously with estrogen receptor antagonists, there were no group differences in object investigation during (**E**) male training. (**F**) In the male testing phase, there was an effect of drug on male investigation time but no effect of ATS (N = 5-8/group, 2-way ANOVA). There were no group differences in object exploration during (**G**) female training. (**H**). In the female testing phase, there was an effect of ATS on investigation, but this effect was not driven by the drug employed (N = 7-11/group, 2-way ANOVA). When estrogen receptor antagonists were infused directly into dorsal hippocampus, there were no group differences in object investigation during (**I**) male training (**J**) male testing (N = 5-9/group, 2-way ANOVA) (**K**) female training (**L**) or female testing (N = 4-7/group, 2-way ANOVA). In the same mice tested two weeks post ATS, there were no group differences in object investigation during (M) male training, (N) male testing, (N = 3-7/group, 2-way ANOVA) (O) female training, (P) or female testing (N = 4-7/group, 2-way ANOVA). Each point represents an individual mouse. Bars display the mean ± SEM.

**Fig. S5.**
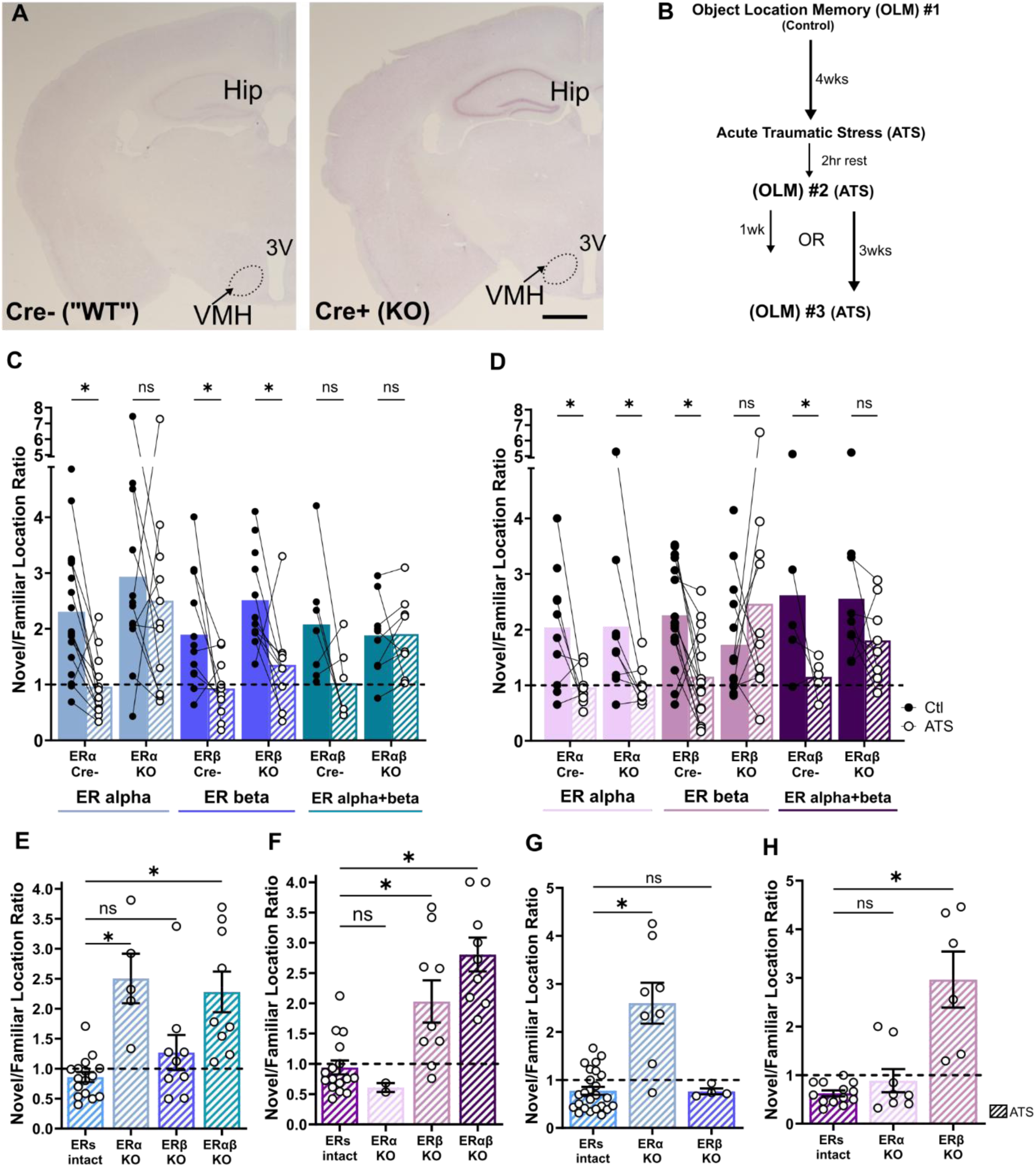
Estrogen receptor intact mice, regardless of floxed ER genotype, are susceptible to lasting disruptions of object location memory. Estrogen receptor knockout enduringly protects memory in a sex-specific manner. (**A**) Cre expression in the transgenic mouse lines employed is almost exclusively in hippocampus (hip). Scale bar = 800 µm. (**B**) Male and female mice of each ERKO genotype were tested on the object location memory task (control). One month later, mice were exposed to ATS (females exposed during proestrus) then OLM was tested again after a 2 hour rest. Approximately half of the mice were tested on a third OLM 1 week after ATS. The remaining mice were tested on a third OLM 3 weeks after ATS. (**C**) ATS disrupts object location memory in all “ER intact” mice (all Cre negative mice) regardless of floxed estrogen receptor genotype. (**D**) ATS disrupts object location in all “ER intact” (all Cre negative mice) proestrous female mice regardless of floxed estrogen receptor genotype. (**E**) One week after ATS, enduring OLM disruptions were observed in ER intact and ERβKO mice, while ERαKO and ErαβKO male mice had intact memory (N = 5-16/group, one-way ANOVA with Sidak’s multiple comparisons). (**F**) One week after ATS, enduring OLM disruptions were observed in ER intact and ERαKO female mice, while ERβKO and ERαβKO male mice had intact memory (N = 2-18/group, one-way ANOVA with Sidak’s multiple comparisons). (**G**) Three weeks after ATS, OLM was impaired in ER intact and ERβKO male mice while ERαKO mice had intact spatial memory (N = 5-25/group, Kruskal Wallis with Dunn’s multiple comparisons). (**H**) Three weeks after ATS, OLM was impaired in ER intact and ERαKO female mice while ERβKO mice had intact spatial memory (N = 6-13/group, Kruskal Wallis with Dunn’s multiple comparisons). Each point represents an individual mouse with connected points representing the same mouse at two time points. Bars display the mean ± SEM. Post-hoc comparisons are displayed above the graph. * = *P*<0.05, # = *P*<0.10

**Fig. S6.**
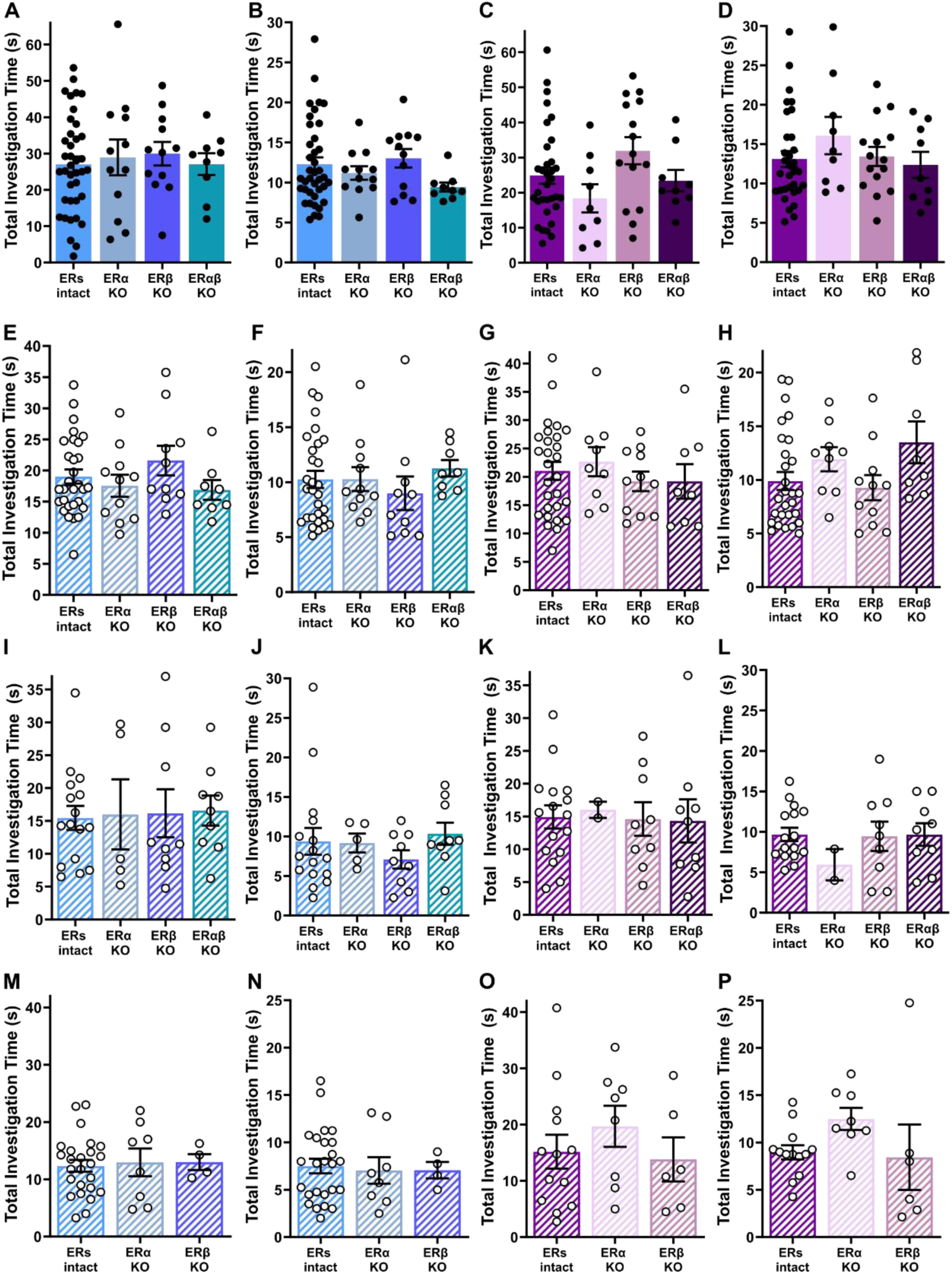
Estrogen receptor knockout protects object location memory from ATS without altering object investigation time. In the control OLM session, there were no group differences in object exploration during (**A**) the male training phase (**B**) the male testing phase (N = 9-39/group, ordinary one-way ANOVA) (**C**) the female training phase, or (**D**) the female testing phase (N = 9-33/group, ordinary one-way ANOVA). In the OLM task conducted two hours after ATS, there were no group differences in object exploration during (**E**) the male training phase (**F**) the male testing phase (N = 8-30/group, ordinary one-way ANOVA) (**G**) the female training phase, or (**H**) the female testing phase (N = 6-24/group, ordinary one-way ANOVA). 1 week after ATS, there were no group differences in object exploration during (**I**) the male training phase, (**J**) the male testing phase (N = 5-17/group, ordinary one-way ANOVA), (**K**) the female training phase, (**L**) or the female testing phase (N= 2-19/group, ordinary one-way ANOVA). 3 weeks after ATS, there were no group differences in object exploration during (**M**) the male training phase, (**N**) the male testing phase (N= 4-25/group, ordinary one-way ANOVA), (**O**) the female training phase, (**P**) or the female testing phase (N= 6-13/group, ordinary one-way ANOVA). Each point represents an individual mouse. Bars display the mean ± SEM.

**Fig. S7.**
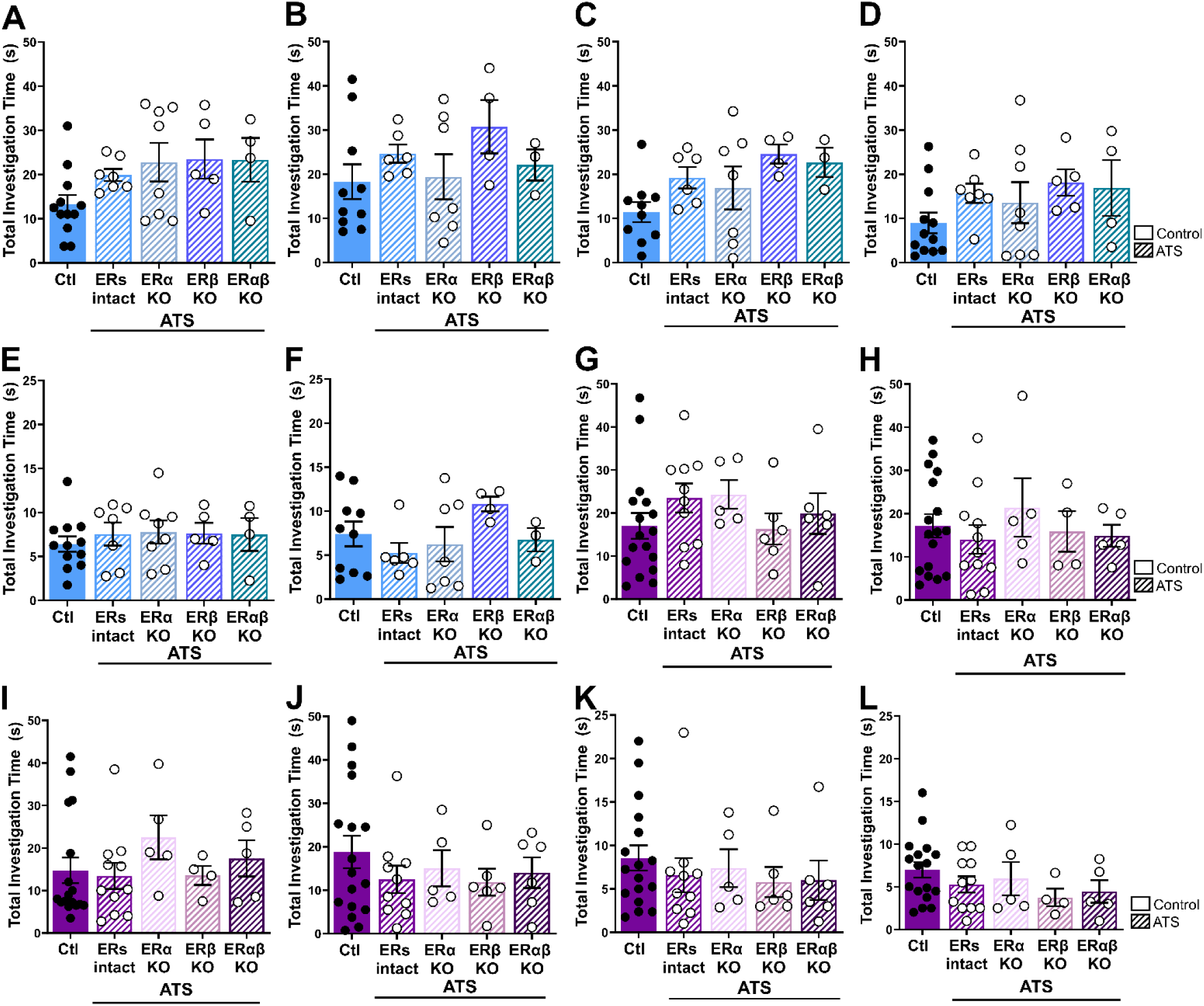
Temporal memory disruptions and protection with estrogen receptor knock out were not a result of differences in investigation times during training or testing. Male and proestrous female ERKO mice were exposed to acute traumatic stress (ATS) and temporal order memory was assessed after a 2hr rest period. In male mice, there were no group differences in object exploration during (**A**) Train A, (**B**) Train B, (**C**) Train C, (**D**) Train D (N = 3-12/group, ordinary one-way ANOVA, Šidák’s multiple comparisons), (**E**) Test A vs D (N = 3-12/group, ordinary one-way ANOVA, Šidák’s multiple comparisons), or (**F**) Test B vs C (N = 3-10/group, ordinary one-way ANOVA, Šidák’s multiple comparisons). In proestrous female mice, there were no group differences in object exploration during (**G**) Train A, (**H**) Train B, (**I**) Train C, (**J**) Train D, (N = 3-17/group, ordinary one-way ANOVA, Šidák’s multiple comparisons), (**K**) Test A vs D (N = 5-17/group, ordinary one-way ANOVA, Šidák’s multiple comparisons), or (**L**) Test B vs C (N = 3-17/group, ordinary one-way ANOVA, Šidák’s multiple comparisons). Each point represents an individual mouse. Bars display the mean ± SEM.

**Fig. S8.**
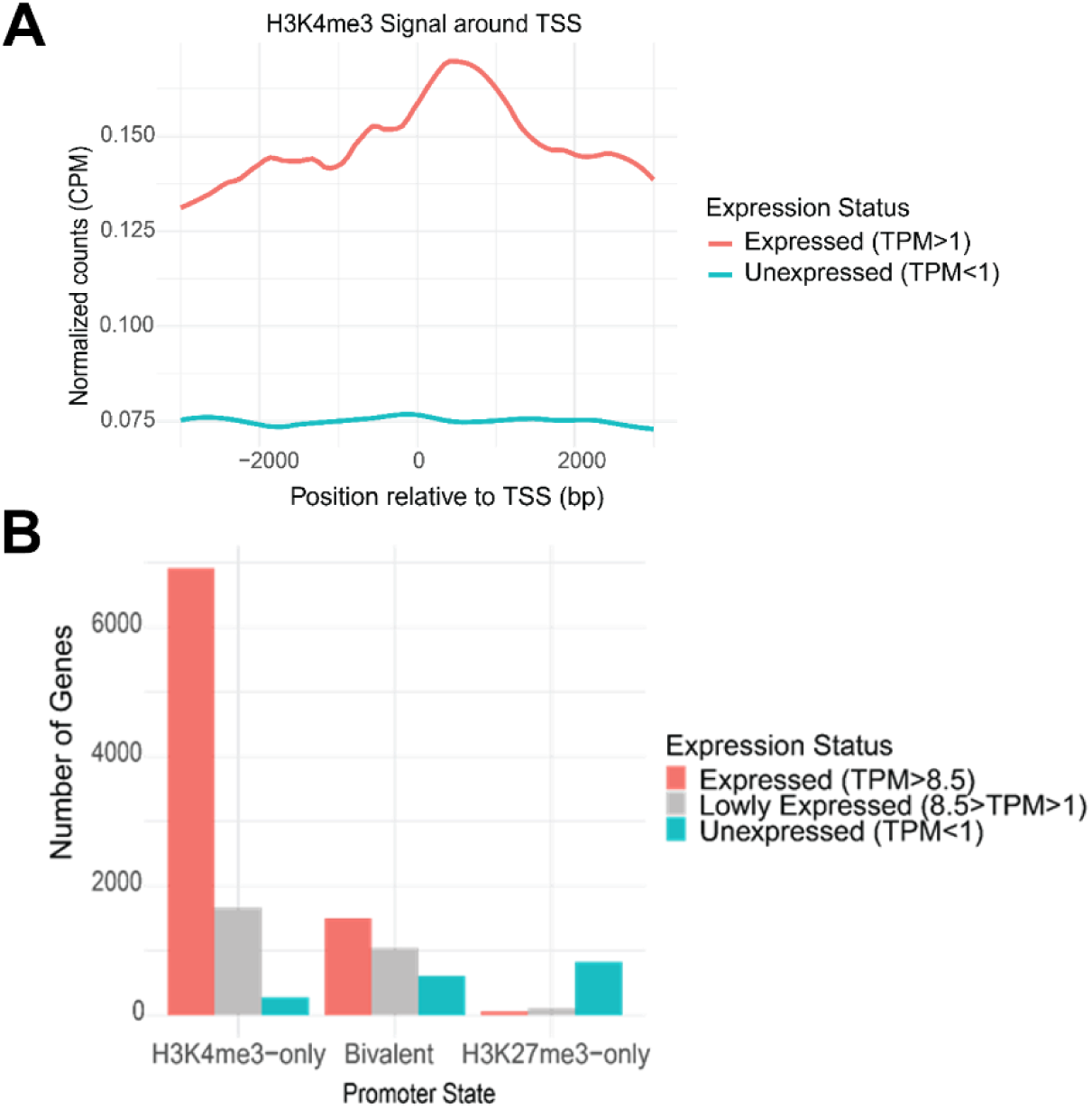
H3K4me3 enrichment correlates with high transcript expression, while H3K27me3 and bivalent chromatin mark lower transcript levels. (**A**) H3K4me3 density near expressed and unexpressed gene promoters. Genes were classified as expressed (TPM > 1) or unexpressed (TPM < 1) based on RNA-seq data. The average H3K4me3 signal across all male, proestrous, and estrous samples is shown. Expressed genes exhibit strong H3K4me3 enrichment near the transcription start site (TSS), while unexpressed genes show minimal signal. (**B**) Expression status of genes by promoter chromatin state. Expressed genes were further classified as lowly expressed (1 < TPM ≤ 8.5, first quartile) or highly expressed (TPM > 8.5). Most H3K4me3-marked promoters are highly expressed, while many bivalent promoters fall into the lowly expressed category. H3K27me3-only promoters are primarily unexpressed.

**Table S1.**
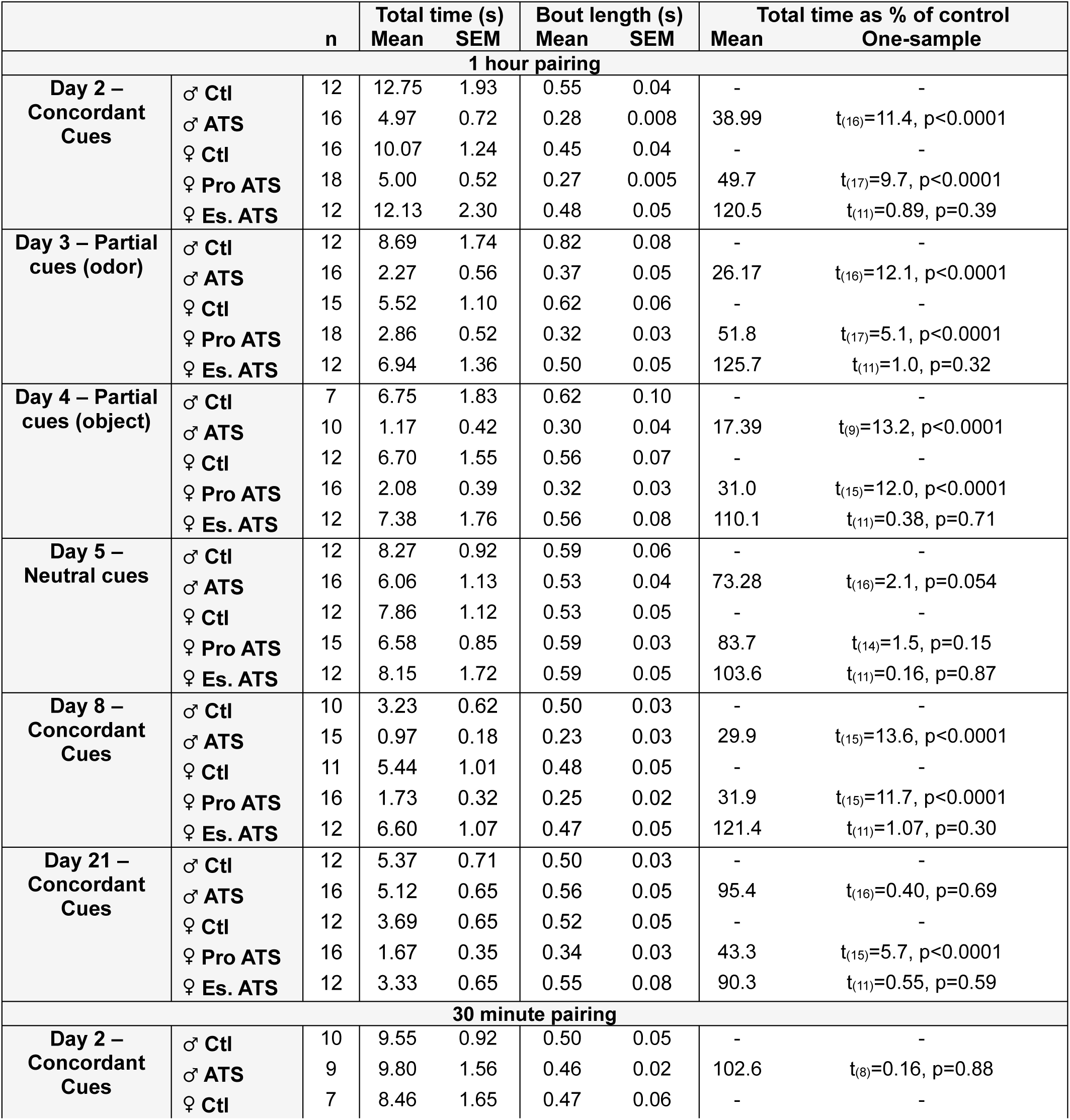
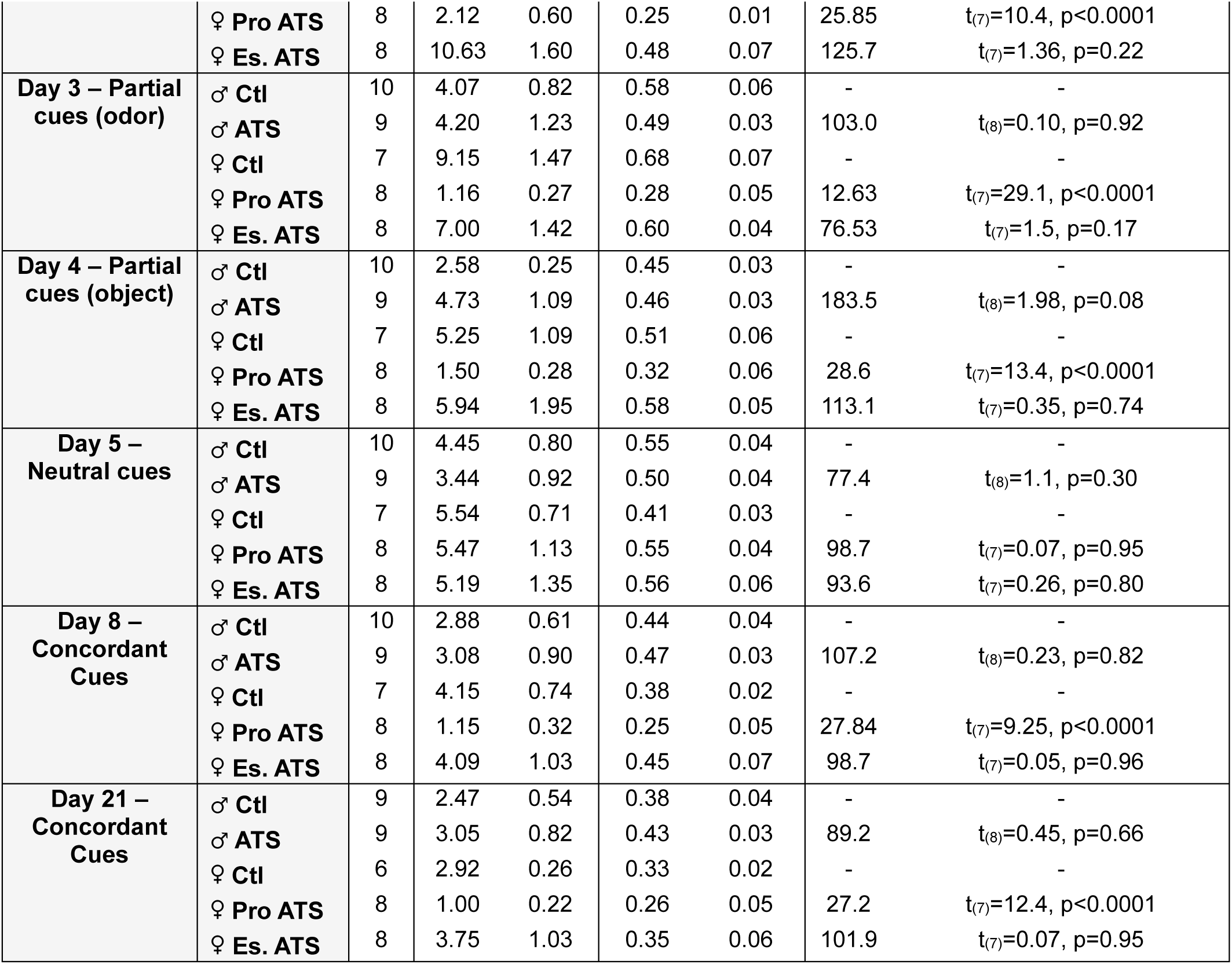
ATS-cue object contacting total time and average bout length in WT males and females. Wild type males and females were exposed to ATS and “concordant cues” (ATS-object and ATS scent) were immediately placed in the home cage for 1 hour or 30 minutes to pair these cues with ATS. For each day, sample size, total object contacting time, and average bout length of object contacting is reported. Total exploration time of ATS groups is also expressed as a % of their same-sex control group. Results of a one-sample T-test (theoretical mean: 100%) determine if exploration time is different than controls.

**Table S2.**
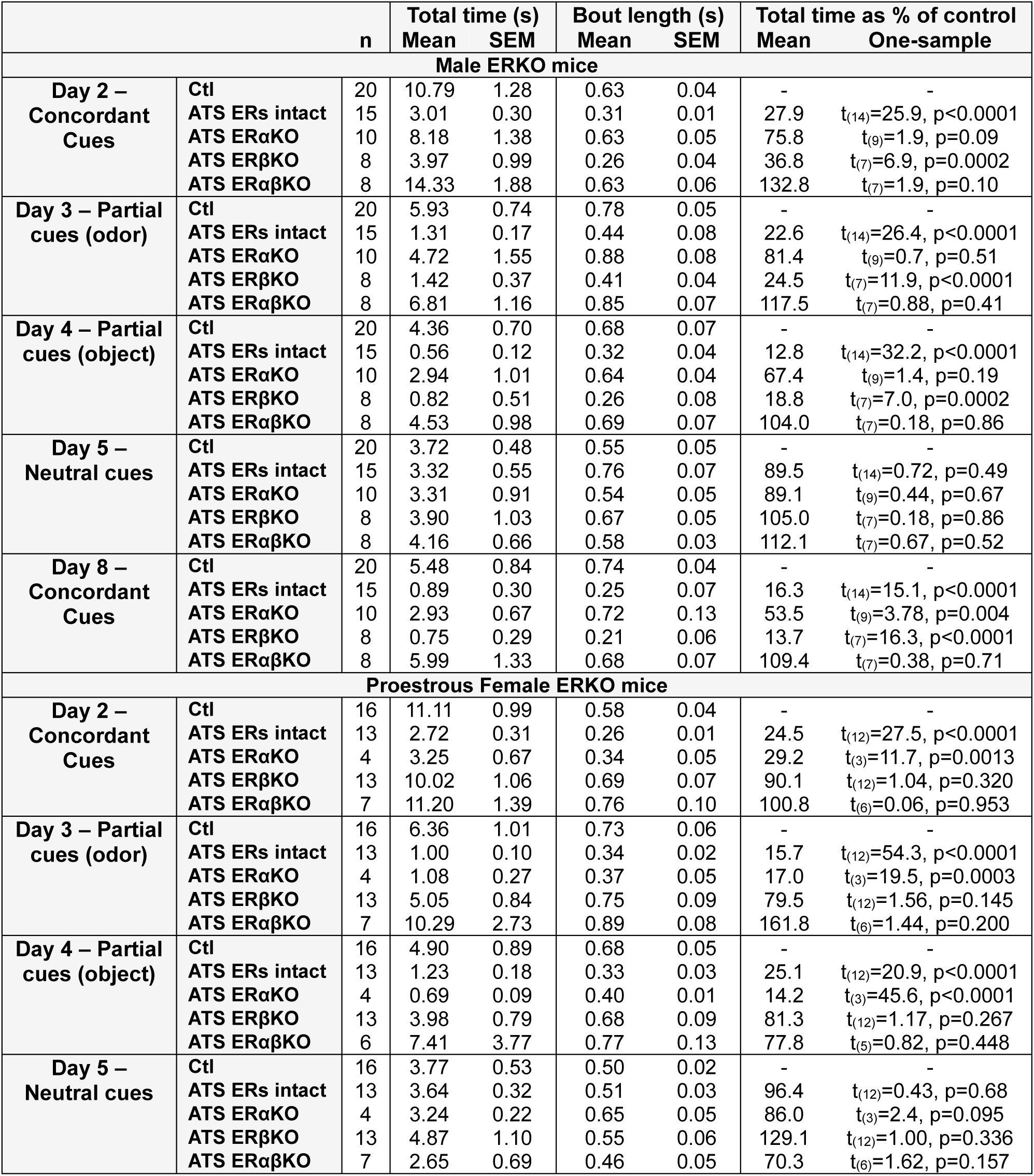
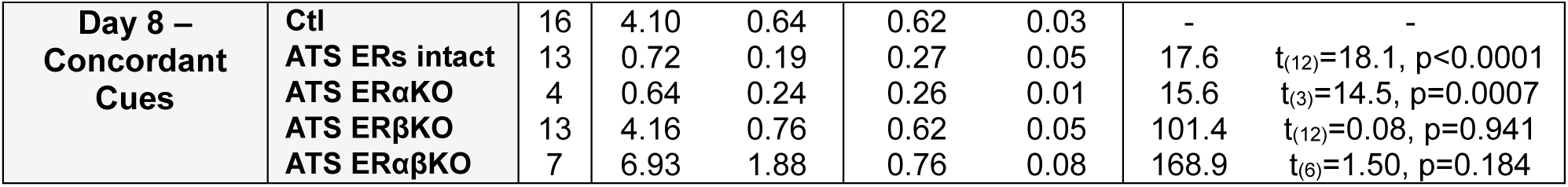
ERKO mice ATS-cue object contacting total time and average bout length. ERKO males and proestrous females were exposed to ATS and “concordant cues” (ATS-object and ATS scent) were immediately placed in the home cage for 1 hour to pair these cues with ATS. For each day, sample size, total object contacting time, and average bout length of object contacting is reported. Total exploration time of ATS groups is also expressed as a % of their same-sex control group. Results of a one-sample T-test (theoretical mean: 100%) determine if exploration time is different than controls.

## Materials and Methods

All experiments were conducted according to the National Institute of Health (NIH) guidelines on laboratory animal welfare and were approved by University of California-Irvine’s Institutional Animal Care and Use Committee (IACUC). Mice were assigned to groups randomly.

### Animals

Two- to six-month old male or female virgin, gonadally intact C57BL6J mice from the Jackson Laboratory or bred in house participated in these studies. Mice were group housed (2-5 mice) by sex in a quiet, uncrowded facility. The housing facility was kept on a 12 hour light cycle (lights on 6:30AM) and mice were given *ad libitum* access to food (Envigo Teklad 2020x global soy protein-free extrude) and water. Mice were housed by stress condition (control cage vs ATS cage) in individually ventilated cages with Envigo 7092-7097 Teklad corncob bedding and iso-BLOX nesting material. The temperature was maintained between 22°C and 24°C.

To investigate the roles of hippocampal estrogen receptors (ERs), ER knock-out transgenic mice (ERKO mice) were generated. We crossed B6;129S6-Tg(Camk2a-cre/ERT2)1Aibs/J (CamK2a-CreERT2) mice (JAX #012362) with estrogen receptor floxed mice: B6(Cg)-Esr1^tm^^4.1Ksk^/J (ERα^lox^) mice, (JAX #032173), C57BL/6N-Esr2^tm1Ksk^/J (Ex3βER^lox^) mice (JAX #032174), or a ERαβ double-floxed mouse generated by crossing the two strains^91–93^. The CamK2a-cre/ERT2 mice express a tamoxifen-inducible Cre-recombinase under the control of the mouse Camk2a promoter. After several generations, mice are homozygous for the floxed ER (α, β, or both α and β) and either carrying cre-recombinase (Cre^+^) or not (Cre^-^). To induce ER deletion, mice were given subcutaneous tamoxifen (10mg/kg/day for 4 days) dissolved in corn oil starting at PND60. ERs were deleted in Cre+ mice and are referred to as “ERαKO”, “ERβKO”, or “ERαβKO”. Mice negative for Cre were also treated with tamoxifen but ERs are not deleted and mice are referred to as “ER intact”. To avoid potential effects of tamoxifen (a mixed ER agonist/antagonist) on behavior, experiments were conducted at least 1 month after tamoxifen administration.

### Estrous Cycle Monitoring

Estrous cycle phases of female mice were monitored daily via vaginal cytology. The mouse was gently lifted, and a phosphate-buffered saline moistened small cotton tipped applicator (Puritan^®^ 890-PC DBL) was inserted into the vagina to scrape for cells. These cells were then smeared across a gelatin-coated microscope slide (Fisherbrand^™^ 12-552-3) and stained using the Shandon^™^ Kwik-Diff^™^ Kit (Thermo Scientific^™^ 9990700). Cell types were identified under a microscope to classify cycle phases. Smears were collected in the morning within four hours after the beginning of the light cycle. On days of ATS (or control), smears were collected as early as two hours prior to the light cycle. Cycles were monitored for a minimum of two complete cycles prior to studies. Mice that were not regularly cycling were not used or experiments were delayed until normal cycling commenced. To ensure that any sex differences are not attributable to handling during estrous cycle monitoring, male mice were handled in a similar manner. For these “sham” smears, the male mouse was gently lifted and the anogenital region of the male mouse was briefly rubbed with the same style cotton swab used in the above vaginal smear protocol.

### Acute Traumatic Stress (ATS)

Male and female mice were assigned to the acute traumatic stress (ATS) or a home-cage (“unstressed”) control group. In the ATS paradigm, mice are exposed to simultaneous physical, emotional, and social stresses^5–7,58^ at the onset of the light phase (6AM). Briefly, mice were individually restrained in a ventilated 50mL plastic tube. Two to ten mice of the same sex were placed in a cage atop a laboratory shaker in a room bathed with loud (90dB) rap music and bright lights (∼400 lux) for two hours. A detailed protocol is available at Bio-protocol^94^. Control mice remained in their home cage and were not exposed to stress. For within-day behavioral assessments, mice underwent ATS for two hours, were returned to the homeroom for one hour, then (alongside control mice). moved to the behavioral testing suite to acclimate for one hour prior to tests. Otherwise, mice were moved to the behavioral testing suite to acclimate for one hour prior to any behavioral testing. Importantly, we have found that recuperated ATS mice explore objects in the OLM and temporal order memory tasks for durations indistinguishable from those of controls^5–7,58^.

### Learning and Memory Tasks Object location memory (OLM)

The hippocampus-dependent object location memory (OLM) task was adapted from^95^ and conducted several hours after ATS or in the weeks following. Mice were handled for two minutes a day for at least six days, in the housing and behavioral suites. After handling, mice were habituated to an empty experimental apparatus for ten minutes a day for about five days. In the training portion of the OLM task (2 hours after the cessation of ATS, or in the days or weeks later), two identical objects were presented to the mouse for ten minutes (or 15 minutes, see Figure 3D-F). Memory tests were conducted 6-24 hours later (inter trial interval identified in each results section, or 4 hours, see figure 3). In the five-minute testing session, one object was displaced while the other remained in the same location (counter-balanced across mice). Object investigation in the training and testing sessions were scored using BORIS version 7^96^ by observers unaware of the experimental conditions. Object “Investigation” was defined as the mouse’s nose being pointed toward the object within 1 cm distance. Object preference was defined as the amount of time exploring the displaced object divided by time exploring the unmoved object, with a ratio of 1 indicating no preference. Total exploration time was calculated and compared across groups.

For enduring impacts on OLM tested 1 week after ATS (Figure 1A-B), female mice were exposed to ATS during proestrus or estrus then trained on the object location memory task one estrous cycle later (i.e. ATS proestrous mice trained in next proestrous phase) or one week later, whichever occurred first. For enduring impacts on OLM tested 4 or 8 weeks after ATS (Figure 3A-F) female mice were exposed to ATS during proestrus or estrus then, regardless of their estrous cycle phase at training, trained on the object location memory task 4 weeks or 8 weeks later. All control female mice make up the control group and ATS groups are divided according to the estrous cycle phase at the time of ATS. The estrous cycle phase at the time of training or testing did not influence results (i.e. ATS proestrous mice had disruptions of OLM regardless of estrous phase at training/test time).

For ERKO mice (Figure 5D-F), object location memory was assessed using a within-subjects design. One month after tamoxifen treatment, mice of each genotype were tested in the object location memory task (OLM control). Mice were trained with two identical objects for 10 minutes. Four hours later, mice were tested with one object displaced for 5 minutes. One month later, all mice were exposed to ATS, with all female mice exposed to ATS during proestrus. Two hours after the cessation of ATS, mice were trained with two different, identical objects for 10 minutes. Four hours later, mice were tested with one object displaced for 5 minutes (OLM ATS). To test lasting memory disruptions or protection with estrogen receptor knockout, approximately half of the mice were tested on a third OLM 1 week later. All other mice were tested on a third OLM 3 weeks later. We found no effect of genotype across Cre negative mice (ERα Cre^-^, ERβ Cre^-^, and ERαβ Cre^-^, Figure S5B-C), thus all Cre negative mice are pooled and referred to as “ER intact” for these studies.

### Temporal Order Memory

The hippocampus-dependent, temporal order memory task was performed as described^97^ and illustrated (Figure 1C). Wild-type mice were handled as described above and habituated to the empty experimental apparatus for 10 minutes a day for 5 days preceding ATS. Following a 2 hour recuperation from ATS, mice were exposed to two identical objects (Object Set A) for 10min (Train A). After a 1h break, mice were exposed to Object Set B (Train B). This procedure is repeated for object sets C and D. To investigate the impacts of ATS on temporal order memory, the mouse’s behavior towards a “remote” versus a more “recent” object was tested. Test A vs D occurs 4h after Train A and tests temporal order memory between objects A and D (separated by 3h). Test B vs C occurs 5h after Train A and tests temporal order memory between objects B and C (separated by 1h). Object investigation in the training and testing sessions were scored using BORIS version 7^96^ by observers unaware of the experimental conditions. Object “Investigation” was defined as the mouse’s nose being pointed toward the object within 1 cm distance. Temporal order memory is expressed as a discrimination index computed through the following equation: [(time with remote object – time with recent object)/(time with remote + time with recent)] x 100.

The temporal order memory task was also conducted in male or proestrous female ERKO mice (Figure 5G-K). ERKO mice were exposed to ATS or served as controls. The temporal order memory task was conducted as described above. The majority of control mice (male and female) are Cre negative mice across all floxed genotypes (ERα Cre^-^, ERβ Cre^-^, and ERαβ Cre^-^) plus some ERαKO, ERβKO, and ERαβKO mice. “ERs intact” ATS mice are Cre negative mice across all floxed genotypes.

### Contextual Reward Learning

In the contextual reward learning task, the time a mouse spends occupying the reward-paired side of a two chambered apparatus is assessed. Mice were acclimated to the palatable food reward (cocoa pebbles) for 3 days prior to testing. In a clean cage, an individual mouse was given free access to 1 gram of cocoa pebbles for 1 hour per day. The day prior to ATS (or control), mice were habituated to the two chambered apparatus for 10 minutes. Chamber occupancy was assessed to determine the mouse’s “preferred” side of the apparatus. The next day, ATS was conducted as described. Two hours after the cessation of ATS, the conditioning sessions were conducted. For the reward conditioning sessions, the mouse was confined to their less-preferred chamber of the apparatus with a 0.5 gram pile of cocoa pebbles for 10 minutes then returned to their home cage. One hour later the mouse was conditioned to the other side (their “preferred” side) of the chamber for 10 minutes without any reward. These conditioning sessions were repeated, such that each mouse received two rewarding and two unrewarded conditioning sessions. Reward and non-reward pairing sessions were randomly counterbalanced across mice, such that the rewarded session was the first session for approximately half of the mice. One hour after the final conditioning session, the mouse returned to the open apparatus for 10 minutes for a test session. Chamber occupancy is assessed using Ethovision Noldus (version 15). Time in the reward paired side is compared between the test session and the habituation session. Palatable food was weighed prior and after acclimation and conditioning sessions.

### ATS-cue memory task

To investigate memory towards cues paired with ATS, a cue memory task was adapted from^98^ and performed as illustrated (Figure 2A). ATS was conducted as described except that a scent (almond) is introduced inside the restraint tube and in the surrounding environment. Immediately after ATS, a restraint tube with almond scent is placed in the home cage of ATS and control mice for 1h to allow association of these “concordant ATS cues” with ATS. Alternatively, a restraint tube with almond scent (approximately 50µL, brand Signature Select) is placed in the home cage of ATS and control mice for only 30 minutes immediately after ATS (with no scent present during ATS). The following day (Day 2), mice are placed in a cage with concordant ATS cues (restraint tube with almond scent) for 5 minutes. On Day 3, mice are tested with partial ATS cues – the odor (a neutral object: bottle, with almond scent) for 5 minutes. On Day 4, mice are tested with partial ATS cues – the object (the restraint tube with a neutral scent: peppermint, brand McCormick) for 5 minutes. On Day 5 mice are placed in a cage with neutral cues (a different neutral object: syringe, with a different neutral scent: orange, brand McCormick) for 5 minutes. Day 8 (1 week following ATS) and Day 21 (3wk later), mice are placed in a cage with concordant ATS cues for 5 minutes. Object contacting was scored using BORIS version 7^96^ by observers unaware of the experimental conditions. Object “contacting” was defined as the mouse’s nose being pointed toward the object within 0.5 cm distance. The mean object contacting time (s) of the control group per sex was set as 100% and data for ATS groups reflects contacting time as a percent of control exploration. Object contacting total times and average bout length are reported in Table S1. ATS and control mice were expected to explore neutral cues equally. Avoidance of the concordant trauma cues is interpreted as a “fear / ATS memory” and avoidance of the partial trauma cues may reflect “generalization” or loss of specificity of this memory. Freezing behavior was not observed in response to cues.

To investigate the role of estrogen receptors in ATS-cue memory, male or proestrous female ERKO mice (Figure 6) were exposed to ATS or served as controls. ATS was conducted as described above, with cues being paired with ATS for 1 hour. The majority of control mice (male and female) are Cre negative mice across all floxed genotypes (ERα Cre^-^, ERβ Cre^-^, ERαβ Cre^-^) plus some ERβKO mice. “ERs intact” ATS mice are Cre negative mice across all floxed genotypes. Object contacting total times and average bout length are reported in Table S2.

### Cannulation surgery

To understand and isolate the role of estrogen receptors in the hippocampus and hippocampus-derived estrogen, mice were implanted with indwelling bilateral dorsal hippocampal cannulae. Mice were anesthetized with an i.p. administration of a 100mg/kg ketamine and 10mg/kg xylazine cocktail diluted in sterile saline or procedures were conducted under isoflurane anesthesia. Mice were implanted with stainless steel bilateral guide cannulae aimed at the dorsal hippocampus (23 gauge hypodermic tubing, -1.9 AP, +/- 1.5 ML, -1.3 DV). Implants were held secure with a stainless-steel anchor screw (Antrin Miniature Specialties, 000-120 x ⅛ SL Bind MS SST or 000-120 x 1/16 SL Pan SST) implanted over the frontal cortex. Plugs, constructed out of 30 gauge stainless steel wire, were inserted into the guide cannulae to prevent clogging. Mice were allowed to recover from surgery for one week before behavioral testing. Mice with cannula implants were individually housed to prevent injury and damage to the cannula.

### Pharmacology

The conversion of androgens to estrogens in hippocampus was blocked with infusing the aromatase inhibitor letrozole into the dorsal hippocampus through indwelling, bilateral cannuale that were implanted at least one week prior (Figure 4A-C). One hour prior to ATS, vehicle (1% DMSO in saline) or letrozole (6ng or 60ng per hemisphere^99^, data were similar and thus pooled for these experiments) were infused at a rate of 0.15 µL/min over 2 min per hemisphere. The infusion cannula remained in the guide cannula for 1 minute post infusion and was replaced with a plug to prevent drug spread up the tract.

Estrogen receptor antagonists were utilized to probe the role of estrogen receptor alpha versus beta in ATS-induced memory disruptions. Mice were given subcutaneous administrations of the ERα antagonist MPP (Methyl-piperidino-pyrazole hydrate, Sigma-Aldrich M7068), the ERβ antagonist PHTPP (4-[2-Phenyl-5,7-*bis*(trifluoromethyl)pyrazolo[1,5-*a*]-pyrimidin-3-yl]phenol, Sigma-Aldrich SML 1355), or corn oil vehicle at 0.5mg/kg 90 minutes before ATS^100^. All subcutaneous treatments were assigned randomly but per cage to avoid cross-contamination. Alternatively, antagonists (MPP at 0.05pmol per hemisphere or PHTPP at 0.1pmol per hemisphere^101,102^) or vehicle (1% DMSO in sterile saline) were infused through indwelling dorsal hippocampal cannulae that were implanted at least one week prior. Compounds were infused at a rate of 0.15 µL/min over 2 min per hemisphere 60 minutes before ATS. The infusion cannulae remained in the guide cannulae for 1 minute post infusion and were replaced with a plug to prevent drug spread up the tract. The intrahippocampal letrozole (Fig4B-C) and intrahippocampal antagonist experiments (Fig4H-I) employ the same mice for the vehicle infused groups.

### Tissue collection for RNA, Chromatin, or Mass Spectrometry Analysis

Mice were euthanized by rapid decapitation. Brains were immediately removed from the skull, hippocampi (bilateral) were dissected on ice (within 2min), flash frozen on dry ice, then stored at -80°C until processing.

### Single sample sequencing (S3EQ)

Single Sample Sequencing (S3EQ) enables exploration of both transcriptomic and epigenomic profiles from the same biological sample. Flash-frozen hippocampal samples were homogenized using a Dounce homogenizer in Cell Lysis Buffer (10 mM Tris pH 8.0, 10 mM NaCl, 3 mM MgCl2, and 0.05% NP-40) and fractionated. The supernatant was collected for mRNA isolation using the Qiagen RNEasy Kit, while the pellet was resuspended for chromatin processing^73^. mRNA was processed with polyA selection and subjected to RNA-sequencing, generating approximately 50 million reads per sample.

### Cleavage Under Targets and Release Using Nuclease (CUT&RUN)

The nuclei pellets were equally divided for incubation with either H3K27me3 antibodies (Active Motif, 39055) or H3K4me3 antibodies (Abcam, Ab8580). Briefly, nuclei were immobilized using 12 µL per reaction of Concanavalin A beads (Epicypher) in Binding buffer (20 mM HEPES pH 7.5, 10 mM KCl, 1 mM CaCl2, 1 mM MnCl2) and Wash buffer (20 mM HEPES pH 7.5, 150 mM NaCl, 0.5 mM spermidine, supplemented with Complete Protease Inhibitor cocktail (Roche). The mixture was rotated for 15 minutes at room temperature. After removing the Wash buffer, the bead-bound nuclei were resuspended in 50 μL of cold Antibody buffer (0.004% digitonin, 2 mM EDTA and 1:50 dilution of antibody) and incubated overnight. The following morning, the antibody buffer was removed and the beads incubated with ProteinA-MNase (Epicypher) for 1 hour at 4°C with rotation. To promote chromatin cleavage by Protein A-MNase, 20 mM CaCl2 was added, and the reaction was incubated for 2 hours at 4°C with rotation. The reaction was terminated with Stop buffer (340 mM NaCl, 20 mM EDTA, 4 mM EGTA, 50 µg/mL RNase A, 200 µg/ML Glycogen) and incubated at 37°C for 10 minutes for chromatin release. DNA was cleaned up with 1 µL of 10% SDS and 1.5 µL of 20 mg/mL Proteinase K followed by purification with the QiaQuick Kit (Qiagen).

### Chromatin Library Preparation

Libraries were prepared using NEBNext® UltraTM II DNA Library Prep Kit for Illumina (New England Biolabs). Libraries were prepared using the NEBNext® Ultra™ II DNA Library Prep Kit for Illumina (New England Biolabs). Adapter sequences were ligated to DNA fragments, which were then amplified by PCR using a Universal i5 primer, uniquely barcoded i7 primers, and NEBNext High-Fidelity 2x PCR Master Mix. The PCR cycle consisted of an initial denaturation at 98°C for 30 seconds, followed by 13 cycles of 98°C for 10 seconds and 65°C for 10 seconds, and a final extension at 65°C for 5 minutes. PCR products were purified using 1.1x volume of Ampure Beads (Beckman Coulter) and washed with 80% ethanol. The cleaned DNA was eluted in 0.1X TE buffer. Library size distribution was confirmed using the Agilent TapeStation System (Agilent Technologies). Libraries were sequenced in paired-end mode on the Illumina HiSeq 2500 platform, generating approximately 20 million reads per sample.

### RNA-Sequencing and Differential Gene Expression Analysis

Raw FASTQ files were processed and quality-checked using FastQC (v0.11.9)^103^. Adapter sequences were trimmed using BBduk (v38.84)^104^. Trimmed reads were pseudoaligned to the mm10 reference genome using Kallisto (v0.46.2)^105^. Kallisto abundance files were imported into RStudio for differential gene expression analysis using DESeq2^106^. Gene ontology analysis was performed using GO-seq^107^ using expressed gene list (TPM>1) as reference set.

### CUT&RUN Sequencing Analysis

Raw FASTQ files were processed and quality-checked using FastQC v0.11.9)^103^. Adapter sequences were trimmed using BBduk (v38.84)^104^. Trimmed reads were aligned to the mm10 reference genome using Bowtie2 (v2.4.1)^108^. Uniquely mapped reads with a quality score >20 were retained using SAMtools (v1.13)^109^. Duplicate reads were removed using the MarkDuplicates command in Picard (v3.0.0)^110^. H3K27me3 and H3K4me3 deduplicated reads were size-selected for fragments between 150–500 bp using SAMtools (v1.13)^109^. Read alignments were normalized to total mapped reads using deepTools (v3.5.4.post1)^111^.

### H3K4me3 and H3K27me3 Bivalency Analysis

ChromHMM (v1.25)^112^ was used to binarize .bam files for H3K4me3 and H3K27me3 into 200 bp bins using the BinarizeBam command with the –paired flag for paired-end reads. Signal from .bam files was averaged between biological replicates for each experimental group. The resulting binary files were used to train the model with the LearnModel command. The whole genome for male, proestrous female, and estrous groups was segmented into four states: H3K4me3-only regions, H3K27me3-only regions, bivalent regions, and non-marked regions. Segmentation files were imported into RStudio along with gene promoter coordinate files (100 bp upstream and downstream from the transcription start site (TSS) for the mm10 mouse genome). Promoters were classified into the four states using the GRanges and FindOverlaps from the GenomicRanges package^113^.

### Chromatin Motif Enrichment Analysis

Motif enrichment analysis was performed using MEME SEA^114^. The JASPAR CORE motif database^115^ was used as the primary reference, and results were validated against other curated transcription factor databases, including Mouse Uniprobe and Jolma. Control sequences were selected from chromatin segments shared across all three groups (estrous, proestrous, and male) for each state (bivalent or permissive). Input sequences were trimmed to the same length to avoid overestimation of significance, as MEME SEA is sensitive to sequence length differences. Significance was assessed using q-values to control for false discovery rate due to multiple hypothesis testing.

### Estrogen analysis by ultrasensitive liquid chromatography tandem mass spectrometry

Estrogens were measured by an ultrasensitive and specific liquid chromatography tandem mass spectrometry (LC-MS/MS) assay in mouse hippocampus. Analytes included estrone (E1), 17β-estradiol (17β-E2), 17α-estradiol (17α-E2), and estriol (E3). Samples were collected from adult female mice during proestrus (N = 8) and estrus (N = 7) and adult male mice (N = 8). Samples were derivatized using 1,2-dimethylimidazole-5-sulfonyl-chloride (DMIS)^116^, which increases sensitivity for estrogens by approximately 10-fold^55^.

Flash frozen hippocampal samples were weighed and placed in Bead Ruptor tubes containing five zirconium ceramic beads (1.4 mm diameter). Samples were homogenized in 1 mL of HPLC-grade acetonitrile at 4 m/s for 30 sec using a bead mill homogenizer. A volume of homogenate equivalent to 2 mg of brain tissue was used for steroid extraction.

Steroids were extracted by liquid-liquid extraction, followed by DMIS derivatization. In brief, we added 50 μL of deuterated internal standard (17β-estradiol-d4; C/D/N Isotopes Inc.) in 50:50 HPLC-grade methanol:MilliQ water to each sample except double blanks to track steroid recovery. Then 1 mL of HPLC-grade acetonitrile was added, samples were vortexed, and samples were centrifuged at 16,100 *g* for 5 min. Then 1 mL of supernatant was transferred to pre-cleaned borosilicate glass culture tubes. Next, 0.5 mL of HPLC-grade hexane was added, and samples were vortexed and then centrifuged at 3,200 *g* for 2 min. The hexane was discarded, and the acetonitrile was dried in a vacuum centrifuge at 60°C for 50 min (ThermoElectron SDP111V; Thermo Fisher Scientific). The dried samples were then placed in a wet ice bath and resuspended in 30 μL of sodium bicarbonate buffer (50 mM, pH 10.5). Then 20 μL of 1 mg/mL DMIS in acetone was added. Samples were vortexed, centrifuged at 3200 *g* for 1 min, and transferred to glass inserts in LC vials. Vials were capped and placed in a water bath at 60°C for 15 min. Samples were cooled at 4°C for 15 min, centrifuged at 3200 *g* for 1 min, and stored at -20°C until injection for LC-MS/MS analysis.

Calibration curves (0.05 – 1000 pg), blanks, double blanks, and quality controls (QCs) were extracted alongside samples. QCs at two different concentrations (2 and 50 pg) were prepared in neat solution and run in triplicate to assess accuracy and precision. Accuracy was assessed by comparing measured values with known values, and precision was assessed by the coefficient of variation (CV) across replicates. Accuracy of all QCs was acceptable (within 15%), and precision was also acceptable (%CV <15%). Analytes were non-detectable in all blanks and double blanks. Underivatized standards were included to measure any underivatized estrogens in samples that underwent derivatization. These standards received 20 μL of acetone without DMIS after resuspension in sodium bicarbonate buffer. In all derivatized standards, blanks, quality controls, and samples, we were unable to detect underivatized estrogens, demonstrating a reaction efficiency of 100%.

Steroids were analyzed using a QTRAP 6500 UHPLC-MS/MS system (Sciex LLC, Framingham MA, USA) as previously described^54,55,117^. Samples were loaded into an autoinjector at 15°C. Then 35 μL of each sample was injected into a Nexera X2 UHPLC system (Shimadzu Corp., Japan). Samples were passed through a KrudKatcher ULTRA HPLC In-Line Filter (Phenomenex, Torrance, CA, USA), then through a SecurityGuard^TM^ ULTRA C18 guard column (2.1 mm) (Phenomenex). Then, samples were separated on a Kinetex® 2.6 µM EVO C18 100 A° LC column (Phenomenex; 2.1 × 50 mm; 2.6 μm; at 40°C) using 0.1 mM ammonium fluoride in MilliQ water as mobile phase A (MPA) and HPLC-grade methanol as mobile phase B (MPB). A gradient elution was used with a flow rate of 0.4 mL/min. During loading, the gradient profile was at 10% MPB for 0.5 min. From 0.6 to 4 min, the gradient profile was increased to 42% MPB, then to 60% MPB until 9.4 min. From 9.4 to 9.5 min, the gradient was at 60-70% MPB and increased to 98% MPB until 11.9 min. Finally, a column wash was carried out from 11.9 to 13.4 min at 98% MPB. The MPB was then returned to the starting conditions of 10% MPB for 1 min. The total run time was 14.9 min. The autoinjector needle was rinsed externally before and after each sample with 100% methanol.

Steroid concentrations were quantified using a 6500 QTRAP triple quadrupole tandem mass spectrometer (Sciex LLC, Framingham MA, USA). Target steroids were detected with two scheduled multiple reaction monitoring (MRM) transitions, and deuterated internal standards were detected with one MRM transition. MRMs were scheduled based on known retention times within a detection window of 3 min. Analytes were ionized using electrospray ionization. Derivatized estrogens were ionized using positive mode, while underivatized estrogens were ionized using negative mode. The lower limit of quantification (LLOQ) was 25pg/g and values under the LLOQ were set to the LLOQ divided by the square root of 2 (17.7pg/g)^118^. The estrogens 17α-estradiol, estrone, and estriol were not detected in any of the hippocampus samples.

### Brain processing and detection of estrogen receptors

Mice were anesthetized with a lethal dose of a 1:10 dilution of Euthasol (488 mg/kg pentobarbital sodium and 63 mg/kg phenytoin sodium, intra-peritoneally) and perfused intracardially with freshly prepared 4% paraformaldehyde in 0.1M sodium phosphate buffer (PB; pH 7.4, 4°C). Brains were cryoprotected and sectioned into 20-30 µm slices. The anti-ER alpha is a rabbit polyclonal antibody from Invitrogen (cat# PA1-309). Brain sections were incubated in the antibody (1:10,000) for 3 days (4°C), followed by incubations in biotinylated anti-rabbit IgG (1:400, 2 h, Vector lab) and avidin-biotin-peroxidase complex (1:200, 3 h, Vector lab) at RT. The reaction product was developed in 3,3’-diaminobenzidine (DAB) containing 0.01% H2O2 (Bioenno Tech) for 10 min. The anti-ER beta is a rabbit polyclonal antibody from ThermoFisher (cat# PA1-310B). Brain sections were incubated in the antibody (1:3,000) for 2-3 days (4°C), followed by standard avidin-biotin-peroxidase immunostaining as described above.

The expression of Cre was detected via in situ hybridization (ISH) as described^119^. Digoxigenin (DIG)−5’-conjugated Cre sense (5’-GGACACAGUGCCCGUGUCGG-3’) and antisense (5’-CCCUUCCAGGGCGCGAGUUG-3’) RNA oligonucleotide probes were generated by GenScript (Piscataway, NJ). The hybridization was performed at 64-66 °C overnight, and hybrid molecules were detected with an anti-DIG serum tagged with alkaline phosphatase (1:1,000, overnight, 4°C) (Roche, #1 093 272). The sections were developed in a mixture of 4-nitro blue tetrazolium chloride (NBT, Roche, #1 383 213) and 5-bromo-4-chloro-3-indolylphosphate (BCIP, Roche, #1 383 221) (25 µl each in 10 ml TBS, pH9.5) for 4-5 h.

### Experimental Design and Statistical Analyses

Data were analyzed using GraphPad Prism version 10.1.2 for Windows (GraphPad software, Inc., La Jolla, CA). As we had found that estrous females and proestrous females are differentially impacted by ATS^7^, when analyzing for sex differences, we used a factor of “sex/cycle”, thus treating male, estrous female, and proestrous female as distinct groups. Statistical tests used to analyze behavior data are as follows: 2-way repeated measures ANOVA was used to analyze trauma cue memory with a 30 minute pairing duration (using factors of ATS group and test day) and contextual reward learning (using factors of sex/cycle and test period). Mixed-effects analysis was used to analyze Wild-type ATS-cue memory (1 hour pairing, using factors of sex/cycle and day), ATS-cue memory in ERKO mice (Figure 6), and ERKO object location memory (control versus ATS, Figure 5, with ATS and genotype as factors). 2-way ANOVA was used to analyze temporal order memory (sex/cycle and ATS as factors), OLM conducted 1 week after ATS, (Figure 1B, sex/cycle and ATS as factors), and antagonist or letrozole treated mice (with ATS and drug as factors). Ordinary one-way ANOVA was used to analyze OLM with 15 minute training sessions (Figure 3E-F), estradiol levels (Figure 4A), temporal order memory in ERKO mice (Figure 5H-K), and 1 week post ATS OLM in ERKO (Figure S5D-E). Data were not normally distributed for 10 minute OLM conducted 4 weeks and 8 weeks after ATS (Figure 3B-C) and 3 weeks post ATS OLM in ERKO mice (Figure S5F-G) and thus was analyzed with Kruskal-Wallis test. For the ATS-cue task, data were additionally analyzed with one-sample t-test, to determine if, per group, ATS mice avoided cues relative to the control group (Table S1 and Table S2). When a main effect or interaction was found to be statistically significant (α = 0.05) or if a specific comparison was planned (cases identified in results), post-tests were run: Sidak’s multiple comparisons for ordinary 1-way ANOVA and 2-way ANOVA, Dunnet’s multiple comparisons for 2-way repeated measures ANOVA and mixed-effects analysis, or Dunn’s multiple comparisons for Kruskal-Wallis tests. Outliers were excluded by ROUT when applicable. Results are reported as mean ± standard error of the mean (SEM) and a data point represents results from one mouse unless noted otherwise.

## Author contributions

REH, KLR, EAH, and TZB designed the research. REH, KLR, YC, AKS, SAS, BD, and BJJ, collected the data. REH, KLR, BJJ, JJW, KKS, EAH, and TZB analyzed data. REH and TZB wrote the original draft of the manuscript. REH, KLR, CMG, KKS, EAH, and TZB edited the manuscript. All authors discussed and commented on the manuscript.

## Acknowledgements

This work was supported by National Institutes of Health: R01 MH 132680 to TZB, P50 MH096889 to TZB, R01 MH126027 to EAH, T32 MH119049-02 to REH, T32 DA050558-03 to REH, the National Science Foundation: NSF GRFP DGE1845298 to KRA, and the Canadian Institutes of Health Research: Project Grant 169203 to KKS, CIHR CGS-M Fellowship to BJJ. We thank Nandini Desaigoudar, Saibrinda Kotthru, Graciella D. Angeles, and Samantha E. Bolotsky for excellent technical assistance.

## Literature Cited

1. Clouston, S. A. P. et al. Cognitive impairment and World Trade Centre-related exposures. Nature Reviews Neurology vol. 18 103–116 Preprint at 10.1038/s41582-021-00576-8 (2022).

2. Lowe, S. R. & Galea, S. The Mental Health Consequences of Mass Shootings. Trauma Violence Abuse 18, 62–82 (2017).

3. Musazzi, L., Tornese, P., Sala, N. & Popoli, M. Acute or Chronic? A Stressful Question. Trends in Neurosciences vol. 40 525–535 Preprint at 10.1016/j.tins.2017.07.002 (2017).

4. Novotney, A. What happens to the survivors. Monitor on Psychology vol. 49 36 (2018).

5. Chen, Y. et al. Converging, synergistic actions of multiple stress hormones mediate enduring memory impairments after acute simultaneous stresses. Journal of Neuroscience 36, 11295– 11307 (2016).

6. Maras, P. M. et al. Preferential loss of dorsal-hippocampus synapses underlies memory impairments provoked by short, multimodal stress. Mol Psychiatry 19, 811–822 (2014).

7. Hokenson, R. E. et al. Unexpected role of physiological estrogen in acute stress-induced memory deficits. The Journal of Neuroscience 41, 648–662 (2021).

8. Chen, Y., Dubé, C. M., Rice, C. J. & Baram, T. Z. Rapid loss of dendritic spines after stress involves derangement of spine dynamics by corticotropin-releasing hormone. Journal of Neuroscience 28, 2903–2911 (2008).

9. Tempesta, D., Mazza, M., Iaria, G., De Gennaro, L. & Ferrara, M. A specific deficit in spatial memory acquisition in post-traumatic stress disorder and the role of sleep in its consolidation. Hippocampus 22, 1154–1163 (2012).

10. Lesuis, S. L. et al. Stress disrupts engram ensembles in lateral amygdala to generalize threat memory in mice. Cell (2024) doi:10.1016/j.cell.2024.10.034.

11. Ressler, K. J. et al. Post-traumatic stress disorder: clinical and translational neuroscience from cells to circuits. Nat Rev Neurol 18, 273–288 (2022).

12. Hunt, C. et al. Pre-deployment threat learning predicts increased risk for post-deployment insomnia: Evidence from the Marine Resiliency Study. Behaviour Research and Therapy 159, (2022).

13. Birkeland, M. S., Blix, I., Solberg, Ø. & Heir, T. Gender differences in posttraumatic stress symptoms after a terrorist attack: A network approach. Front Psychol 8, (2017).

14. Carmassi, C. et al. DSM-5 PTSD and posttraumatic stress spectrum in Italian emergency personnel: Correlations with work and social adjustment. Neuropsychiatr Dis Treat 12, 375–381 (2016).

15. Fullerton, C. S. et al. Gender Differences in Posttraumatic Stress Disorder After Motor Vehicle Accidents. Am J Psychiatry 158, 1486–1491 (2001).

16. Solomon, Z., Gelkopf, M. & Bleich, A. Is terror gender-blind? Gender differences in reaction to terror events. Soc Psychiatry Psychiatr Epidemiol 40, 947–954 (2005).

17. Sever, I., Somer, E., Ruvio, A. & Soref, E. Gender, distress, and coping in response to terrorism. Affilia - Journal of Women and Social Work 23, 156–166 (2008).

18. Woolley, C. S. & McEwen, B. S. Roles of estradiol and progesterone in regulation of hippocampal dendritic spine density during the estrous cycle in the rat. Journal of Comparative Neurology 336, 293–306 (1993).

19. Goldstein, J. M. et al. Normal Sexual Dimorphism of the Adult Human Brain Assessed by In Vivo Magnetic Resonance Imaging. Cerebral Cortex 11, 490–497 (2001).

20. Pritschet, L. et al. Functional reorganization of brain networks across the human menstrual cycle. Neuroimage 220, (2020).

21. Rocks, D., Cham, H. & Kundakovic, M. Why the estrous cycle matters for neuroscience. Biol Sex Differ 1–14 (2022) doi:10.1186/s13293-022-00466-8.

22. Bale, T. L. & Epperson, C. N. Sex differences and stress across the lifespan. Nat Neurosci 18, 1413–1420 (2015).

23. Becker, J. B. & Chartoff, E. Sex differences in neural mechanisms mediating reward and addiction. Neuropsychopharmacology vol. 44 166–183 Preprint at 10.1038/s41386-018-0125-6 (2019).

24. Gegenhuber, B., Wu, M. V., Bronstein, R. & Tollkuhn, J. Gene regulation by gonadal hormone receptors underlies brain sex differences. Nature 606, 153–159 (2022).

25. Noel, X. & Christiansen, D. M. Editorial: Understanding the influences of sex and gender differences in mental disorders. Front Psychiatry 13, (2022).

26. Gall, C. M., Le, A. A. & Lynch, G. Sex differences in synaptic plasticity underlying learning. J Neurosci Res 1–19 (2021) doi:10.1002/jnr.24844.

27. Bowman, R. E., Zrull, M. C. & Luine, V. N. Chronic restraint stress enhances radial arm maze performance in female rats. Brain Res 904, 279–289 (2001).

28. Liu, J. et al. Acute Restraint Stress Increases Intrahypothalamic Oestradiol Concentrations in Conjunction with Increased Hypothalamic Oestrogen Receptor β and Aromatase mRNA Expression in Female Rats. J Neuroendocrinol 23, 435–443 (2011).

29. Ortiz, J. B. et al. Sex-specific impairment and recovery of spatial learning following the end of chronic unpredictable restraint stress: Potential relevance of limbic GAD. Behavioural Brain Research 282, 176–184 (2015).

30. Heck, A. L. & Handa, R. J. Sex differences in the hypothalamic–pituitary–adrenal axis’ response to stress: an important role for gonadal hormones. Neuropsychopharmacology vol. 44 45–58 Preprint at 10.1038/s41386-018-0167-9 (2019).

31. Peay, D. N. et al. Chronic unpredictable intermittent restraint stress disrupts spatial memory in male, but not female rats. Behavioural Brain Research 383, (2020).

32. Zuloaga, D. G., Heck, A. L., De Guzman, R. M. & Handa, R. J. Roles for androgens in mediating the sex differences of neuroendocrine and behavioral stress responses. Biol Sex Differ 11, 1–18 (2020).

33. Tan, T. et al. Neural circuits and activity dynamics underlying sex-specific effects of chronic social isolation stress. Cell Rep 34, 108874 (2021).

34. Kato, A. et al. Female hippocampal estrogens have a significant correlation with cyclic fluctuation of hippocampal spines. Front Neural Circuits 7, 1–13 (2013).

35. Hojo, Y. & Kawato, S. Neurosteroids in adult hippocampus of male and female rodents: Biosynthesis and actions of sex steroids. Frontiers in Endocrinology Preprint at 10.3389/fendo.2018.00183 (2018).

36. Nilsson, M. E. et al. Measurement of a comprehensive sex steroid profile in rodent serum by high-sensitive gas chromatography-tandem mass spectrometry. Endocrinology 156, 2492–2502 (2015).

37. Jaric, I., Rocks, D., Greally, J. M., Suzuki, M. & Kundakovic, M. Chromatin organization in the female mouse brain fluctuates across the oestrous cycle. Nat Commun 10, (2019).

38. Vierk, R. et al. Aromatase inhibition abolishes LTP generation in female but not in male mice. Journal of Neuroscience 32, 8116–8126 (2012).

39. Frick, K. M., Kim, J., Tuscher, J. J. & Fortress, A. M. Sex steroid hormones matter for learning and memory: Estrogenic regulation of hippocampal function in male and female rodents. Learning and Memory 22, 472–493 (2015).

40. Oberlander, J. G. & Woolley, C. S. 17β-Estradiol acutely potentiates glutamatergic synaptic transmission in the hippocampus through distinct mechanisms in males and females. Journal of Neuroscience 37, 12314–12327 (2017).

41. Wang, W. et al. Memory-related synaptic plasticity is sexually dimorphic in rodent hippocampus. Journal of Neuroscience 38, 7935–7951 (2018).

42. Lu, Y. et al. Neuron-derived estrogen regulates synaptic plasticity and memory. Journal of Neuroscience 39, 2792–2809 (2019).

43. Taxier, L. R., Gross, K. S. & Frick, K. M. Oestradiol as a neuromodulator of learning and memory. Nat Rev Neurosci 21, 535–550 (2020).

44. Luine, V. & Frankfurt, M. Estrogenic regulation of memory: The first 50 years. Hormones and Behavior vol. 121 Preprint at 10.1016/j.yhbeh.2020.104711 (2020).

45. Conrad, C. D., McLaughlin, K. J., Huynh, T. N., El-Ashmawy, M. & Sparks, M. Chronic stress and a cyclic regimen of estradiol administration separately facilitate spatial memory: Relationship with hippocampal CA1 spine density and dendritic complexity. Behavioral Neuroscience 126, 142– 156 (2012).

46. Rincón-Cortés, M., Herman, J. P., Lupien, S., Maguire, J. & Shansky, R. M. Stress: Influence of sex, reproductive status and gender. Neurobiol Stress 10, (2019).

47. Doncheck, E. M. et al. Estradiol regulation of the prelimbic cortex and the reinstatement of cocaine seeking in female rats. Journal of Neuroscience 41, 5303–5314 (2021).

48. Gore, I. R. & Gould, E. Developmental and adult stress: effects of steroids and neurosteroids. Stress (Amsterdam, Netherlands) vol. 27 2317856 Preprint at 10.1080/10253890.2024.2317856 (2024).

49. Wei, J. et al. Estrogen protects against the detrimental effects of repeated stress on glutamatergic transmission and cognition. Mol Psychiatry 19, 588–598 (2014).

50. Azcoitia, I., Barreto, G. E. & Garcia-Segura, L. M. Molecular mechanisms and cellular events involved in the neuroprotective actions of estradiol. Analysis of sex differences. Front Neuroendocrinol 55, 100787 (2019).

51. Shors, T. J., Lewczyk, C., Pacynski, M., Mathew, P. R. & Pickett, J. Stages of estrous mediate the stress-induced impairment of associative learning in the female rat. Neuroreport 9, 419–423 (1998).

52. Wood, G. E., Beylin, A. V & Shors, T. J. The Contribution of Adrenal and Reproductive Hormones to the Opposing Effects of Stress on Trace Conditioning in Males Versus Females. 115, 175–187 (2001).

53. Cioffi, L. et al. Neuroactive steroids fluctuate with regional specificity in the central and peripheral nervous system across the rat estrous cycle. Journal of Steroid Biochemistry and Molecular Biology 243, (2024).

54. Jalabert, C., Ma, C. & Soma, K. K. Profiling of systemic and brain steroids in male songbirds: Seasonal changes in neurosteroids. J Neuroendocrinol 33, 1–16 (2021).

55. Jalabert, C. et al. Ultrasensitive quantification of multiple estrogens in songbird blood and microdissected brain by LC-MS / MS. eNeuro 9, 1–16 (2022).

56. Tabatadze, N., Sato, S. M. & Woolley, C. S. Quantitative analysis of long-form aromatase mRNA in the male and female rat brain. PLoS One 9, (2014).

57. Fester, L. et al. Control of aromatase in hippocampal neurons. Journal of Steroid Biochemistry and Molecular Biology 160, 9–14 (2016).

58. Hokenson, R. E. et al. Sex-dependent effects of multiple acute concurrent stresses on memory: a role for hippocampal estrogens. Front Behav Neurosci 16, 1–12 (2022).

59. Fuentes, N. & Silveyra, P. Estrogen receptor signaling mechanisms. in Advances in Protein Chemistry and Structural Biology vol. 116 135–170 (Elsevier Inc., 2019).

60. Fu, X. D. & Simoncini, T. Extra-nuclear signaling of estrogen receptors. IUBMB Life 60, 502–510 (2008).

61. Correa, S. M. et al. An estrogen-responsive module in the ventromedial hypothalamus selectively drives sex-specific activity in females. Cell Rep 10, 62–74 (2015).

62. Ingraham, H. A., Herber, C. B. & Krause, W. C. Running the Female Power Grid Across Lifespan Through Brain Estrogen Signaling. Annu Rev Physiol 84, 59–85 (2022).

63. Levin, E. R. & Hammes, S. R. Nuclear receptors outside the nucleus: extranuclear signalling by steroid receptors. Nat Rev Mol Cell Biol 17, 783–797 (2016).

64. Guertin, M. J. et al. Targeted H3R26 Deimination Specifically Facilitates Estrogen Receptor Binding by Modifying Nucleosome Structure. PLoS Genet 10, (2014).

65. Magnani, L. & Lupien, M. Chromatin and epigenetic determinants of estrogen receptor alpha (ESR1) signaling. Mol Cell Endocrinol 382, 633–641 (2014).

66. Day, J. J. & Sweatt, J. D. Epigenetic Mechanisms in Cognition. Neuron 70, 813–829 (2011).

67. Alberini, C. M. & Kandel, E. R. The regulation of transcription in memory consolidation. Cold Spring Harb Perspect Biol 7, (2015).

68. Yap, E. L. & Greenberg, M. E. Activity-Regulated Transcription: Bridging the Gap between Neural Activity and Behavior. Neuron 100, 330–348 (2018).

69. Campbell, R. R. & Wood, M. A. How the epigenome integrates information and reshapes the synapse. Nature Reviews Neuroscience vol. 20 133–147 Preprint at (2019).

70. Brito, D. V. C., Kupke, J., Karaca, K. G., Zeuch, B. & Oliveira, A. M. M. Mimicking age-associated Gadd45γ dysregulation results in memory impairments in young adult mice. Journal of Neuroscience 40, 1197–1210 (2020).

71. Woodfield, G. W., Hitchler, M. J., Chen, Y., Domann, F. E. & Weigel, R. J. Interaction of TFAP2C with the estrogen receptor-α promoter is controlled by chromatin structure. Clinical Cancer Research 15, 3672–3679 (2009).

72. Bernstein, B. E. et al. A Bivalent Chromatin Structure Marks Key Developmental Genes in Embryonic Stem Cells. Cell 125, 315–326 (2006).

73. Xu, S. J. & Heller, E. A. Single sample sequencing (S3EQ) of epigenome and transcriptome in nucleus accumbens. J Neurosci Methods 308, 62–73 (2018).

74. Barha, C. K., Dalton, G. L. & Galea, L. A. M. Low doses of 17α-estradiol and 17β-estradiol facilitate, whereas higher doses of estrone and 17α-and 17β-estradiol impair, contextual fear conditioning in adult female rats. Neuropsychopharmacology 35, 547–559 (2010).

75. Diamond, D. M., Campbell, A. M., Park, C. R., Halonen, J. & Zoladz, P. R. The temporal dynamics model of emotional memory processing: A synthesis on the neurobiological basis of stress-induced amnesia, flashbulb and traumatic memories, and the Yerkes-Dodson law. Neural Plasticity vol. 2007 Preprint at 10.1155/2007/60803 (2007).

76. Joëls, M. & Baram, T. Z. The neuro-symphony of stress. Nat Rev Neurosci 10, 459–466 (2009).

77. Salehi, B., Cordero, M. I. & Sandi, C. Learning under stress: The inverted-U-shape function revisited. Learning and Memory 17, 522–530 (2010).

78. Smiley, C. E. et al. Estrogen Receptor Beta in the Central Amygdala Regulates the Deleterious Behavioral and Neuronal Consequences of Repeated Social Stress in Female Rats. Neurobiol Stress 23, 100531 (2022).

79. Carter, J. S., Costa, C. C., Kearns, A. M. & Reichel, C. M. Inhibition of Estradiol Signaling in the Basolateral Amygdala Impairs Extinction Memory Recall for Heroin-Conditioned Cues in a Sex-Specific Manner. Neuroendocrinology 114, 207–222 (2024).

80. Brailoiu, E. et al. Distribution and characterization of estrogen receptor G protein-coupled receptor 30 in the rat central nervous system. Journal of Endocrinology 193, 311–321 (2007).

81. Lalmansingh, A. S. & Uht, R. M. Estradiol regulates corticotropin-releasing hormone gene (crh) expression in a rapid and phasic manner that parallels estrogen receptor-α and -β recruitment to a 3′,5′-cyclic adenosine 5′-monophosphate regulatory region of the proximal crh promoter. Endocrinology 149, 346–357 (2008).

82. Qi, Y. J. et al. Rapid membrane effect of estrogens on stimulation of corticotropin-releasing hormone. Psychoneuroendocrinology 117, 104680 (2020).

83. Bangasser, D. A. & Wiersielis, K. R. Sex differences in stress responses: a critical role for corticotropin-releasing factor. Hormones Preprint at 10.1007/s42000-018-0002-z (2018).

84. Ivy, A. S. et al. Hippocampal dysfunction and cognitive impairments provoked by chronic early-life stress involve excessive activation of CRH receptors. Journal of Neuroscience 30, 13005– 13015 (2010).

85. Fenoglio, K. A. et al. Enduring, handling-evoked enhancement of hippocampal memory function and GR expression involves activation of the CRF type-1 receptor. Endocrinology 146, 4090– 4096 (2005).

86. Millan, M. J. et al. Cognitive dysfunction in psychiatric disorders: Characteristics, causes and the quest for improved therapy. Nat Rev Drug Discov 11, 141–168 (2012).

87. Christiansen, D. M. & Hansen, M. Accounting for sex differences in PTSD: A multi-variable mediation model. Eur J Psychotraumatol 6, 1–10 (2015).

88. Olff, M. Sex and gender differences in post-traumatic stress disorder: an update. Eur J Psychotraumatol 8, 1351204 (2017).

89. Garcia, N. M., Walker, R. S. & Zoellner, L. A. Estrogen, progesterone, and the menstrual cycle: A systematic review of fear learning, intrusive memories, and PTSD. Clin Psychol Rev 66, 80–96 (2018).

90. Jacobs, E. G. Bridging the neuroscience gender divide. Nature 623, 667 (2023).

91. Binder, A. K. et al. The absence of ER-β results in altered gene expression in ovarian granulosa cells isolated from in vivo preovulatory follicles. Endocrinology 154, 2174–2187 (2013).

92. Hewitt, S. C. et al. Biological and biochemical consequences of global deletion of exon 3 from the ERα gene. The FASEB Journal 24, 4660–4667 (2010).

93. Madisen, L. et al. A robust and high-throughput Cre reporting and characterization system for the whole mouse brain. Nat Neurosci 13, 133–140 (2010).

94. Hokenson, R. E. et al. Multiple Simultaneous Acute Stresses in Mice: Single or Repeated Induction. Bio Protoc 10, e3699 (2020).

95. Vogel-Ciernia, A. & Wood, M. A. Examining object location and object recognition memory in mice. Curr Protoc Neurosci 2014, 8.31.1-8.31.17 (2014).

96. Friard, O. & Gamba, M. BORIS: a free, versatile open-source event-logging software for video/audio coding and live observations. Methods Ecol Evol 7, 1325–1330 (2016).

97. Barker, G. R. I. et al. Separate elements of episodic memory subserved by distinct hippocampal-prefrontal connections. Nat Neurosci 20, 242–250 (2017).

98. Park, S., Cho, J. & Huh, Y. Role of the anterior insular cortex in restraint-stress induced fear behaviors. Sci Rep 12, 1–12 (2022).

99. Tuscher, J. J. et al. Inhibition of local estrogen synthesis in the hippocampus impairs hippocampal memory consolidation in ovariectomized female mice. Horm Behav 83, 60–67 (2016).

100. Amiresmaili, S. et al. The Hepatoprotective mechanisms of 17β-estradiol after traumatic brain injury in male rats: Classical and non-classical estrogen receptors. Ecotoxicol Environ Saf 213, 111987 (2021).

101. Compton, D. R. et al. Pyrazolo[1,5-α]pyrimidines: Estrogen receptor ligands possessing estrogen receptor β antagonist activity. J Med Chem 47, 5872–5893 (2004).

102. Zhou, H. B., Carlson, K. E., Stossi, F., Katzenellenbogen, B. S. & Katzenellenbogen, J. A. Analogs of methyl-piperidinopyrazole (MPP): Antiestrogens with estrogen receptor α selective activity. Bioorg Med Chem Lett 19, 108–110 (2009).

103. Andrews, S. FastQC: A Quality Control Tool for High Throughput Sequence Data. Preprint at https://www.bioinformatics.babraham.ac.uk/projects/fastqc/ (2010).

104. Bushnell, B. BB Tools. Preprint at https://sourceforge.net/projects/bbmap/ (2014).

105. Bray, N. L., Pimentel, H., Melsted, P. & Pachter, L. Near-optimal probabilistic RNA-seq quantification. Nat Biotechnol 34, 525–527 (2016).

106. Love, M. I., Huber, W. & Anders, S. Moderated estimation of fold change and dispersion for RNA-seq data with DESeq2. Genome Biol 15, (2014).

107. Young, M. D., Wakefield, M. J., Smyth, G. K. & Oshlack, A. Open Access METHOD Gene Ontology Analysis for RNA-Seq: Accounting for Selection Bias GOseq GOseq Is a Method for GO Analysis of RNA-Seq Data That Takes into Account the Length Bias Inherent in RNA-Seq. Genome Biology vol. 11 http://genomebiology.com/2010/11/2/R14 (2010).

108. Langmead, B. & Salzberg, S. L. Fast gapped-read alignment with Bowtie 2. Nat Methods 9, 357– 359 (2012).

109. Li, H. et al. The Sequence Alignment/Map format and SAMtools. Bioinformatics 25, 2078–2079 (2009).

110. Broad Institute. Picard Tools. Preprint at http://broadinstitute.github.io/picard/ (2019).

111. Ramírez, F., Dündar, F., Diehl, S., Grüning, B. A. & Manke, T. DeepTools: A flexible platform for exploring deep-sequencing data. Nucleic Acids Res 42, (2014).

112. Ernst, J. & Kellis, M. ChromHMM: Automating chromatin-state discovery and characterization. Nature Methods vol. 9 215–216 Preprint at 10.1038/nmeth.1906 (2012).

113. Lawrence, M. et al. Software for Computing and Annotating Genomic Ranges. PLoS Comput Biol 9, (2013).

114. Bailey, T. L. & Grant, C. E. SEA: Simple Enrichment Analysis of motifs. bioRxiv (2021) doi:10.1101/2021.08.23.457422.

115. Castro-Mondragon, J. A. et al. JASPAR 2022: The 9th release of the open-access database of transcription factor binding profiles. Nucleic Acids Res 50, D165–D173 (2022).

116. Handelsman, D. J. et al. Ultrasensitive serum estradiol measurement by liquid chromatography-mass spectrometry in postmenopausal women and mice. J Endocr Soc 4, 1–12 (2020).

117. Hamden, J. E. et al. Steroid profiling of glucocorticoids in microdissected mouse brain across development. Dev Neurobiol 81, 189–206 (2021).

118. Handelsman, D. J. & Ly, L. P. An Accurate Substitution Method to Minimize Left Censoring Bias in Serum Steroid Measurements. Endocrinology 160, 2395–2400 (2019).

119. Birnie, M. T. et al. Stress-induced plasticity of a CRH/GABA projection disrupts reward behaviors in mice. Nat Commun 14, (2023).

